# An absence of temporal filling-in at the physiological blind-spot

**DOI:** 10.1101/2020.10.13.338285

**Authors:** Neha Dhupia, D. Samuel Schwarzkopf, Derek H. Arnold

**Author notes:** Corresponding Author: Neha Dhupia.

## Abstract

Visual objects that extend across physiological blind spots seem to encapsulate the extent of blindness, due to a process commonly referred to as a perceptual filling-in of spatial vision. It is unclear if temporal perception is similar, so we examined temporal relationships governing causality perception across the blind spot. We found the human brain does not allow for the time an object should take to traverse the blind-spot when engaging in a causal interaction. We also used electroencephalogram (EEG), to examine temporal signatures of elements flickering on and off in tandem, or in counter-phase. At a control site, we found more brain activity was entrained at the duty cycle by flicker relative to counter-phase changes, whereas these conditions were indistinguishable about blind spots. Our data suggest a common pool of neurons might encode temporal properties on either side of physiological blind-spots. This would explain the absence of any allowance for the extent of blindness in causality perception, and the weakened differences between temporal representations of flicker and counter-phased changes about the blind spot. Overall, our data suggest that, unlike spatial vision, there is no temporal filling-in for perceptual representations about physiological blind spots.

## Introduction

The physiological blind spot refers to a region of blindness all sighted people have in each eye, corresponding with the region of retina known as the optic disc [1]. This is the location where the optic nerve passes through the surface of the retina to carry signals to the brain, which dictates that there can be no photoreceptors to absorb light [2]. These regions are quite large. In width they correspond to approximately half the width of the image of your hand when you look at it held upright at an arms’ length [1, 3]. Yet people are often unaware they have blind spots. The biggest reason for this lack of awareness is that the physiological blind spot in each eye encodes images from *different* regions, on opposite sides of your visual field. So if both eyes are open, each eye can compensate for the region of blindness in the other eye. However, even when we close one eye we do not become aware of our temporary blindness to a large region of the visual field.

Another reason why we are often unaware of the blindness associated with the physiological blind spot is because human vision generates experiences that complete across these regions [4, 5]. Importantly, these representations encapsulate the extent of blindness associated with the physiological blind spot without any apparent disruption [6–9]. In spatial vision the processes that result in an allowance for the extent of blindness resulting from the physiological blind spot are commonly referred to as perceptual filling-in [5, 6]. As yet, it is unclear if temporal perception makes similar allowances, and it is this possibility that we turn to investigating here.

To the best of our knowledge, nobody has yet asked if temporal perception is *directly* analogous to spatial vision about the physiological blind spot, in terms of perceptual filling-in. One study did examine how far moving objects seem to complete *into* a blind spot, finding evidence that this may be modulated by motion signals [10]. However, this investigation suffered from some uncertainty, due to the probabilistic nature of vision in regions abutting the blind spot. Not only can small involuntary eye movements shift regions of blindness associated with the physiological blind spot from trial to trial [11], but photosensitivity does not cut off as a step function as you transition into the optic disc, but rather it degrades over short spatial interval [12]. This creates uncertainty when trying to index the extent of a visual experience relative to the edge of a blind spot, as estimates of the blind spot’s edge can only ever be approximate. Consequently, representations that seem to extend a small amount *into* a blind spot might just reflect a more successful recruitment of the limited photosensitivity abutting the blind spot than some other task [13]. For these reasons, we shall investigate a situation that is more analogous to situations that trigger a spatial filling in of perception across the blind spot – we shall examine temporal interactions between elements that are *clearly visible* at opposite edges of a physiological blind spot.

In Experiment 1, we examine a contact-launch scenario, wherein an initially moving object seems to stop when it strikes an initially static object, launching the latter object into motion. This protocol has become a standard paradigm for investigating computations underlying causality perception [14–16], and here we extend its use to investigate causality perception across the physiological blind spot. We find no evidence for an allowance of the extent of blindness when visual objects seem to engage in a causal interaction across the blind spot. In Experiment 2 we use electroencephalogram (EEG) to examine the temporal signatures of elements that flicker on and off in tandem, or in counter-phase to create a sense of apparent motion. These signatures are reasonably well-matched at the blind spot, but differ at a control site matched for eccentricity, eye of origin and side of the visual field. In combination, we feel our data suggest that temporal properties on either side of physiological blind-spots can be encoded by a common pool of neurons, with overlapping receptive fields – explaining the absence of any allowance for the extent of blindness in causality perception, and the weakened differences between temporal representations of flicker and apparent motion across the blind spot. Initial analysis procedures for experiment 1 were predetermined and preregistered (https://aspredicted.org/blind.php?x=my5pu9), experiment 2 was not preregistered.

## Experiment 1: Causality Perception

### Methods

#### Participants

Forty-nine participants (24 males & 25 females), age between 20-35 yrs., volunteered to participate. All had normal or corrected to normal visual acuity (e.g. they were asked to wear corrective lenses if they need to for reading), were naïve as to the purpose of the study, and they were awarded course credit for participation. All participants provided informed consent to participate, and were free to withdraw from the study at any time without penalty. This investigation was granted ethical approval from the University of Queensland Ethics Committee, and was carried out in accordance with the Code of Ethics of the World Medical Association (Declaration of Helsinki).

#### Apparatus

Stimuli were presented on a 19-in Samsung SyncMaster 900SL monitor, or on a Sony Tco99 monitor. Both test displays had a resolution of 1280 x 1024 pixels and a refresh rate of 75Hz. Stimuli were generated using Psychophysics Toolbox for Matlab [17, 18]. Participants viewed stimuli from a distance of 57 cm, using a chinrest to stabilize their head, while wearing a patch over their *left* eye (in order to target investigations at the physiological blind spot in the *right* eye).

#### Stimuli and procedure

All trials, in calibration and experimental tasks, began with a sequence that ensured the participant was fixated on a point on the upper left side of the test display. This began with a red dot, with a diameter subtending 0.6 degrees of visual angle (dva) at the retina, appearing 0.4 dva directly above or below (determined at random on a trial-by-trial basis) a red ring (diameter 0.5 dva). Participant were instructed to fixate this dot throughout the trial. To trigger the beginning of a trial, participants had to use a mouse to ‘drag’ this red dot into the ring. As soon as the dot entered the ring, it turned green and froze in place – at the centre of the ring. As soon as this happened, the trial commenced.

#### Preliminary blind spot localisation tasks

We used a sequence of preliminary tasks to estimate the location, width and height of the physiological blind spot for each participant. First, we had people report when slow (2.3 dva / sec) moving dots disappeared, because they had moved into the blind spot, first from the left, then right, then from above, and finally then from below (in the latter two cases, the moving dots were centred on the x-screen coordinate suggested by reported left and right disappearances). Then we had people report when slow moving dots *appeared*, due to moving out from the middle of the initial estimate of the blind spot location. In a final calibration task, two green bars (vertically centered on the blind spot) appeared to the left and right edges of the blind spot. These were initially seen as clearly separated, and participants were asked to stretch and shorten them, by pressing the right and left mouse buttons respectively, until they seemed to *just* join, forming a solid perceptual bar across the blind spot (although they were physically separated – located on either side of the blind spot). Code for all calibration procedures is available as supplemental material (will be made available via UQeSpace as well).

Our calibration procedures provided an estimate of each participants’ blind spot position (M eccentricity 15.9 dva, SD 1.15) width (M 2.4 dva, SD 1.17) and height (M 2.0 dva, SD 1.04).

#### Causality perception across the blind spot

Test trials began with a green bar (width 1.0 dva, height 0.5 dva), vertically centered on the blind spot, which moved from 8.5 dva to the *left* of the blindspot toward the blindspot at a speed of 1.6 dva/second for 5 seconds. The green bar stopped with its right edge *abutting* the left edge of the blind spot (see Figure 1a). At the start of each trial a larger static red square (width / height 1.0 dva,) was positioned with its’ left edge abutting the *right* side of the blind spot. This initially static red square would begin moving, to the right at a speed of 1.6 dva / second, after some delay. The precise delay was adjusted according to the method of constant stimuli, from 1 second *before* the green bar reached the left edge of the blind spot, to 1 second *after* the green bar reached the left edge – in steps of 0.25 seconds. After each trial participants reported if the red bar had seemed to start moving *before* it was ‘hit’ by the green bar (too early responses), *after* it was hit by the green bar (too late responses), or if it had seemed to start moving as soon as it was ‘hit’ by the green bar.

**Figure 1.**
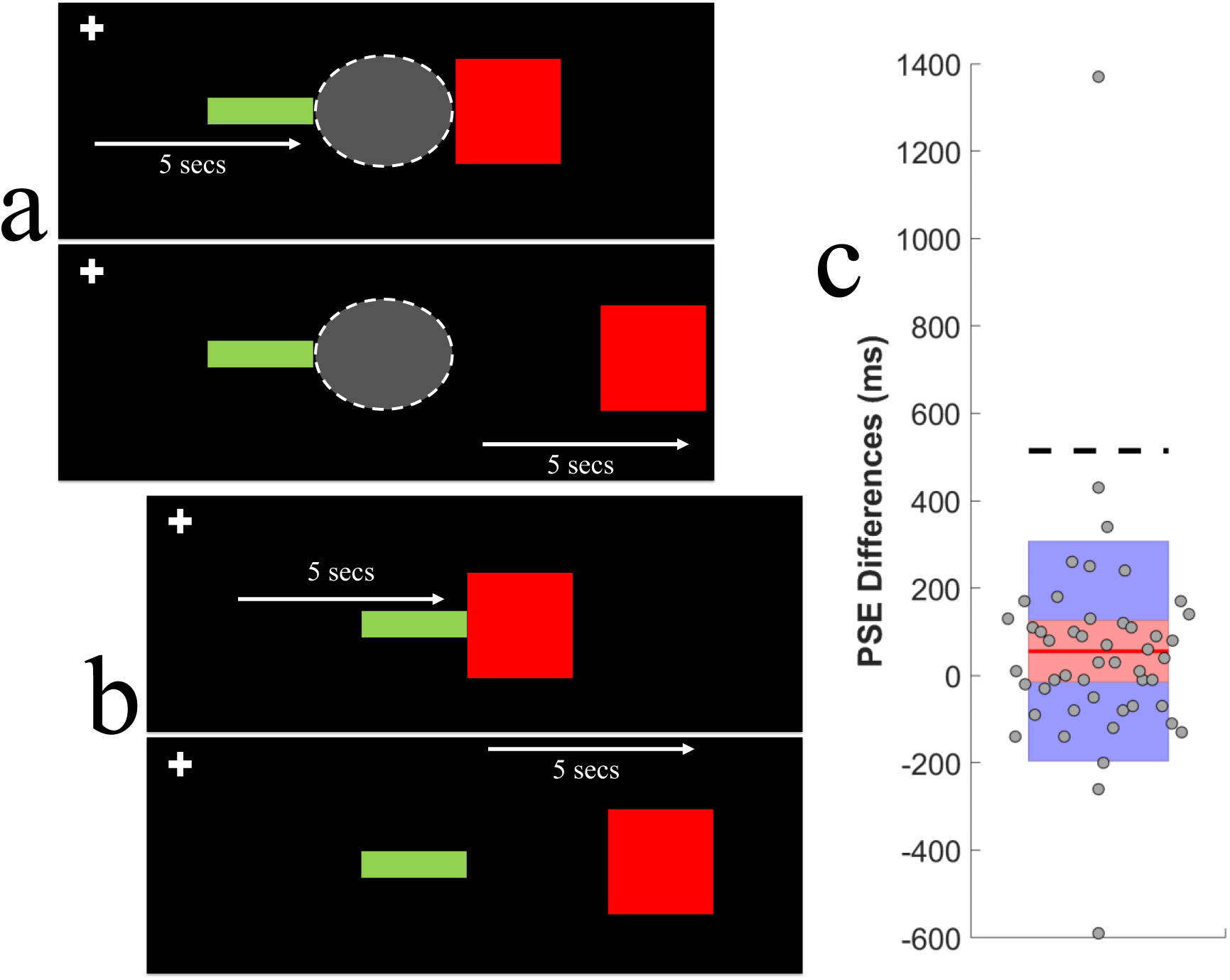
a) Graphic depicting the Causality Perception test protocol, centred on the right eyes’ physiological blind spot. The blind spot is depicted as a shaded oval, with a dotted white outline (this was not visible, the background was a uniform black). ***(above)*** An initially moving green bar would translate for 5 seconds, stopping at the nasal (left) side of the blind spot. ***(below)*** A red square would begin moving at variable times (+/− 1sec) relative to this moment. **b)** As for Figure 1a, but for tests presented at the control position (rotated 60 angular degrees clockwise from the centre of the persons’ physiological blind spot). When the red square started moving after, or as soon as, the green bar stopped, the two bars would physically abut. **c)** Differences in optimal times for motion onset, for the red square to have seemed launched into motion by a collision with the green bar (blind spot tests - control tests). Red line representing the blind spot and shaded red regions represent both sides of the blind spot site whereas shaded blue regions are representing the control site. Individual data points are randomly scattered about the X axis. The difference in time required for the green bar to translate ***half way*** across the blind spot is depicted by the bold dotted horizontal line.

Details for control trials were as for test trials, except that test positions were rotated 60° clockwise relative to the estimated centre of each participants’ blind spot, and the gap corresponding with the width of the blind spot was eliminated – such that the right side of the green bar would physically abut the left side of the red square on trials when the red square started moving as soon as the green bar stopped moving, or after this had happened (see Figure 1b).

A block of trials consisted of 162 individual trials, with 9 red square motion onset delays each sampled 9 times for each type of test (blind spot and control site). These were all completed in random order during a single block of trials. Sigmoid functions were fit to individual data describing ‘too early’ and ‘too late’ responses, as a function of red square motion onset delays – separately for each condition. The 50% points of these two fitted functions were taken as estimates of the temporal threshold for deciding the red square had begun moving too early or too late (respectively) to have seemed to be launched into motion by the green bar. We took midpoints between these two estimates as the individual’s *optimal* motion onset delay – for the red square to have seemed to be ‘launched’ into motion by the green bar. Each block of trials provided two estimates for each participant, one for tests at the blind spot, and one for tests at the control site.

On 25% of trials we also asked participants if the green bar had seemed to accelerate or change shape *before* it had stopped moving. We intermittently asked this question to ensure we had accurately estimated the position of the left edge of the blind spot. If the green bar had entered the blind spot, it might have seemed to suddenly stretch into the blind spot, or accelerate across to touch the red bar on the far side. By intermittently asking these questions in test and control trials, we can determine if there was any apparent distortion of shape (or change in speed) *specific* to the blind spot.

### Results

If casual perception allows for the distance an initially moving object should traverse, in order to cross the physiological blind spot to contact a target on the other side, we would expect this to be reflected in *when* the target should begin moving (to seem to have been launched). We found no evidence for this. The optimal time for target motion onset, to create an impression of a launch, was not measurably delayed at the blind spot (see Figure 1c), relative to a control site (where there was no blindness and the interacting objects could physically ‘touch’, see Figure 1b; M delay 56ms, SD 251; t_48_ = 1.55, p = 0.12; see Figure 1c). This timing was greatly different relative to the time the initially moving target would need to traverse even half-way across the blind spot (M 514ms SD 200; t_48_ = 13.0, p < 0.001; see Figure 1c). Time for stimulus to reach halfway to the blind spot was calculated using ratio of the half width of the blind spot for each individual and the speed at which the stimulus was traversing.

An allowance for the time it would take to traverse the blind spot would be unnecessary if, when the initially moving object reached the blind spot it seemed to either suddenly accelerate, or to deform and stretch across the blind spot. For these reasons we regularly asked participants to report if either of these scenarios had happened. This was only reported on a small proportion of trials when this question was asked (M 0.27 SD 0.28), and not more often relative to a control site where there was no blindness (M 0.26 SD 0.28; t_48_ = 0.09, p = 0.93). From these results it would seem that any reports of changes in the shape or speed of objects was due to visual ambiguity in the periphery of vision, rather than to operations specifically targeting inputs about the blind spot.

The results of Experiment 1 suggest that temporal perception is unlike spatial vision, in that no allowance seems to be made for the extent of blindness associated with the physiological blind spot when objects are perceived as engaging in a causal relationship predicated on timing. On the basis of these data, there would seem to be no temporal filling-in at the blind spot. One plausible explanation of these data would be if temporal dynamics on either side of a physiological blind spot were encoded, at least in part, by a common pool of neurons. This would weaken the power of activity entrained by rhythmic visual stimulation at the blind spot, relative to spatially and temporally matching stimulation at a different site, as stimulations at the blind spot would be encoded by a smaller overlapping pool of neurons. We decided to examine this possibility in Experiment 2.

## Experiment 2: Temporal Sensitivity

### Method

#### Participants

Twenty-one participants (13 males & 8 females), age between 20-35 yrs., volunteered to participate. All had normal or corrected to normal vision (e.g. they were asked to wear corrective lenses if they needed to for reading), were naïve as to the purpose of the study, and were awarded course credit for participation. All participants provided informed consent to participate, and were free to withdraw from the study at any time without penalty. This investigation was granted ethical approval from the University of Queensland Ethics Committee, and was carried out in accordance with the Code of Ethics of the World Medical Association (Declaration of Helsinki).

#### Apparatus

Stimuli were generated using a ViSaGe stimulus generator from Cambridge Research Systems, driven by MATLAB R2015b software, and displayed on an Asus Monitor (resolution 1920 x 1080, refresh rate 60 Hz), viewed from distance of 57cm, with the participants chin restrained by a chin rest. A Biosemi International ActiveTwo system was used to record brain activity. A standard 64 AG/agCI electrode placement was used, according to the extended international 10-20 system. Data were digitised at a 1024Hz sample rate with 24-bit analog-digital conversion. As in Experiment 1, participants wore an eye patch over their left eye, so we could target visual stimulation about the physiological blind spot of the right eye.

### Stimuli and procedure

#### Preliminary blind spot localisation tasks

Participants completed the same sequence of preliminary calibration tasks as Experiment 1, to estimate the location and extent of the physiological blind spot in their right eye. On average, these were located at an eccentricity of 17.3 dva into the right visual field (SD 1.03), and they subtended 4 dva in width (SD 0.85) and 3.1 dva in height (SD 1.88).

#### Measuring the temporal dynamics of brain activity

Test animations consisted of two white square elements (width / height subtending 0.5 dva) which were each flashed on (125ms) and off (125ms) against a black background at a duty cycle of 4 Hz for 10 seconds per trial (see Figure 2a and 2b). The two elements either flashed on and off in tandem (to generate a sense of flicker) or in counter phase (potentially generating a sense of apparent motion). The two elements either abutted the blind spot edges (i.e. the right edge of left element abutted the left edge of the blind spot, and the left edge of the right element abutted the right edge of the blind spot), or they were presented centered on a control site, rotated 60° clockwise from the blind spot and separated by the same physical distance (i.e. the width of the blind spot). As an attentional check, participants were asked after each trial if the elements had been flashing in tandem, or in counter phase. Feedback was provided on this judgment to maintain motivation levels.

**Figure 2.**
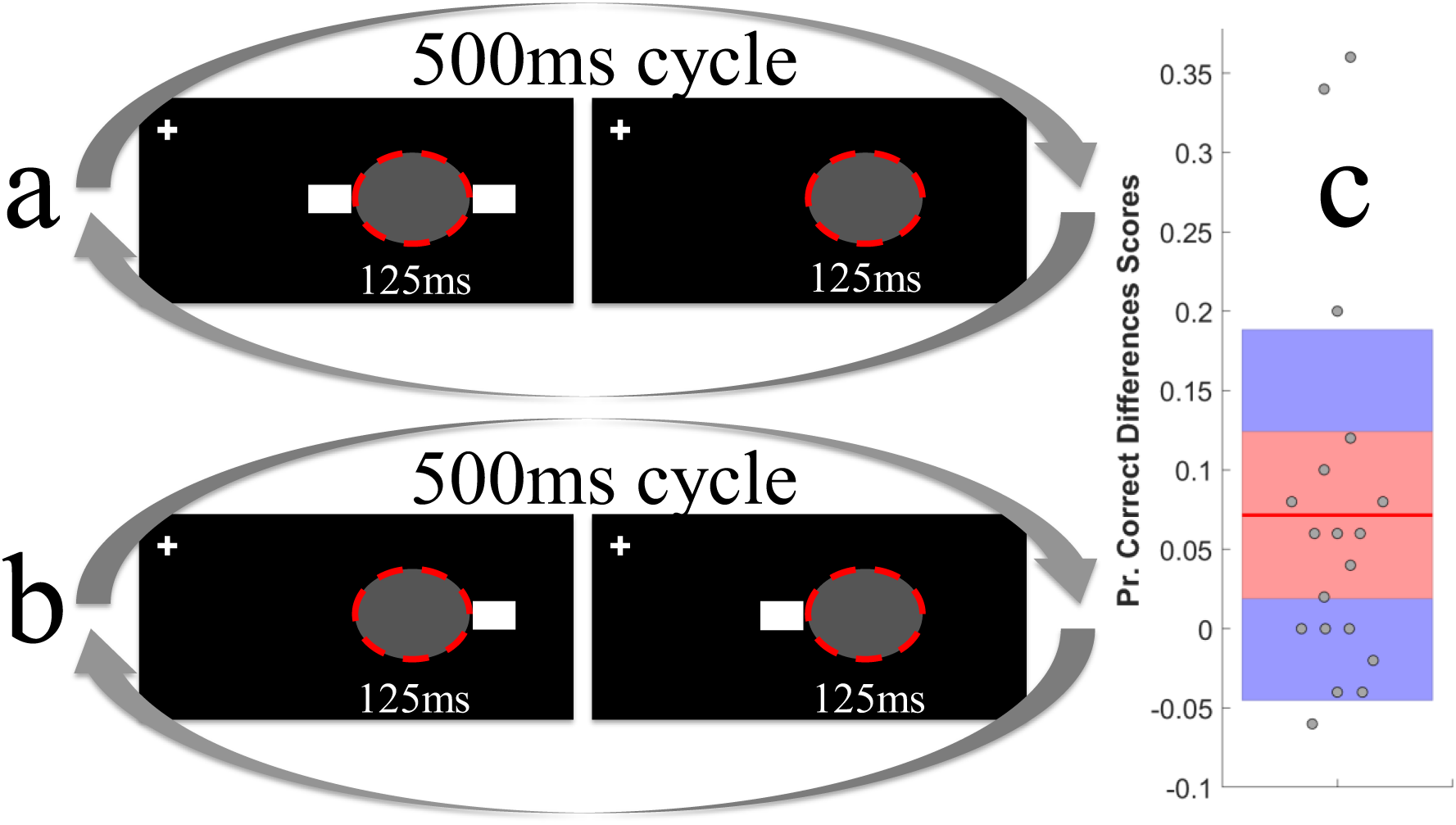
a) Graphic depicting the flicker test condition, in the temporal sensitivity task. **b)** Graphic depicting the apparent motion test condition, in the temporal sensitivity task. **c)** Difference in proportions of correct temporal judgments (flicker vs apparent motion) for blind spot and control site tests, where the shaded red region represent the difference at blind spot site and blue regions indicating the control site.

A block of trials contained 50 blind spot and 50 control site trials, all presented in a pseudo random order in a single block of trials. EEG data analyses were performed using custom MATLAB scripts using the FieldTrip toolbox [19]. EEG data were re-referenced to the average activity of all 64 electrodes, and low (40 Hz) and high (1 Hz) pass filtered. Data were then subjected to a principal-components analysis, to remove blink and movement related artefacts. Trials were then sorted into blind spot and control site presentations. Time-frequency representations (TFRs) of power were calculated using a sliding-window Fourier analysis with a Hanning taper and a time window of 1 second.

### Results

Our attentional check revealed that people had minimal difficulty discerning flicker from apparent motion presentations, at either the control site (M difference 0.045, SD 0.061; t_18_ = 3.34, p = 0.003) or at the blind spot (M difference −0.013, SD 0.075; t_18_ = 0.77, p = 0.453), although people did make slightly more errors when tests were presented at the blind spot (M 7% difference, SD 12; t_18_ = 2.67, p = 0.016; see Figure 2c).

We conducted three non-parametric cluster-based permutation test procedures, based on paired t-tests and implemented using the Fieldtrip toolbox (www.filedtriptoolbox.org) for Matlab [19]. These allow for analyses of conditional differences in 4Hz power that encompass all sensors, while controlling for type 1 error rate. The first involved brain activity evoked by blind spot tests, the second involved brain activity evoked by control site tests, and the third involved a contrast of activity evoked by tests at these two sites. All tests related to differences in evoked 4Hz power, the first two contrasted flicker and apparent motion tests, the third contrasted flicker tests at the control site and at the blind spot. In each test samples with an uncorrected p-value < 0.05 were clustered based on spatio-temporal proximity, independently for positive and negative test results. Cluster-level statistics were then obtained by summing statistics within each cluster, and taking the maximum to test significance against a random distribution, obtained via 1000 permutations of the original data.

We found we could reject the null hypothesis (that there was no difference) for control site flicker vs apparent motion trials (see Figure 3a), and for control site flicker vs blind spot flicker trials (see Figure 3c), but not for blind spot flicker vs apparent motion (see Figure 3b). These data suggest that more brain activity was entrained by coherent flicker at the control site, relative to the blind spot. Moreover, they suggest that the amount of entrained activity was more equally matched across test conditions at the blind spot, relative to the control site.

**Figure 3.**
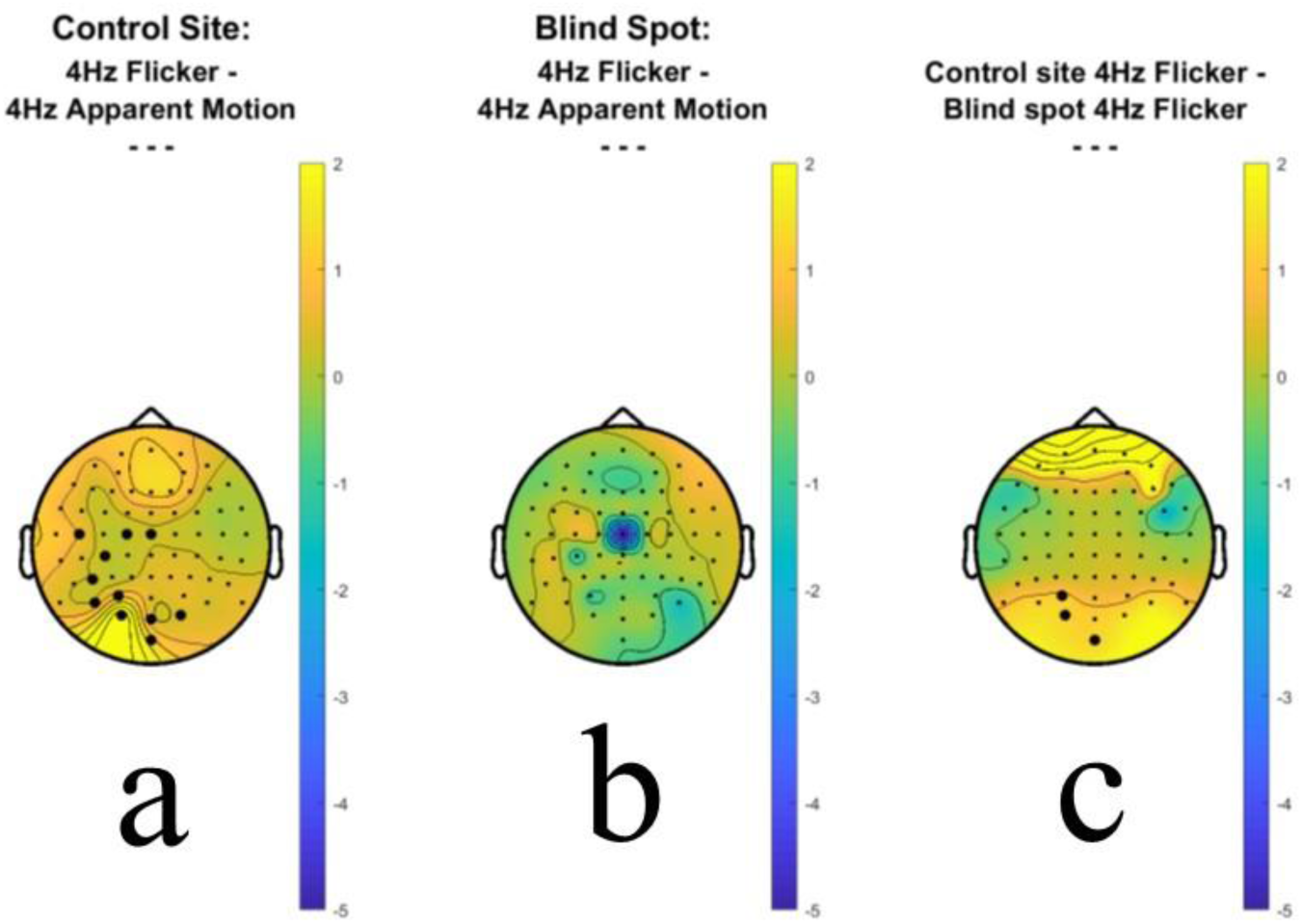
a) Heat map of differences in 4Hz power during flicker relative to apparent motion test presentations at the control test site. Sensors contributing to a positive cluster test, marking sites of enhanced 4Hz power, are highlighted via large black circles. **b)** Heat map of differences in 4 Hz power during flicker relative to apparent motion test presentations at the blind spot. **c)** Heat map of differences in 4Hz power during flicker at the Control test site relative to flicker at the blind spot. Sensors contributing to a positive cluster test, marking sites of enhanced 4Hz power for control site presentations, are highlighted via large black circles.

## General Discussion

We have found that when elements seem to engage in a causal relationship [14, 15] across a physiological blind spot, no allowance is made for the extent of blindness. We have also observed that there is a lesser difference in the amount of brain activity entrained by elements flashing in tandem, relative to temporally offset presentations, when presentations are located on either side of a physiological blind spot, relative to a control site matched for visual eccentricity and visual hemi-field. Also, our data suggest that more brain activity overall tends to be entrained by flashing elements at sites other than those immediately abutting the blind spot. Each of these observations are consistent with dynamics on either side of a physiological blind spot being encoded, at least in part, by a common pool of neurons.

In Experiment 2 our data were suggestive of a lateralisation that is consistent with the hemi-field locus of test presentations. We observed clusters of enhanced 4Hz power during flicker presentations at the control site (relative to apparent motion tests at the control site; see Figure 3a), and during flicker presentations at the control site (relative to flicker presentations at the blind spot; see Figure 3b). Each of these clusters extended anteriorly from a maximal point of difference corresponding with the left occipital sensor 01. These lateralisations are consistent with the right visual field locus of test presentations.

Our data are suggestive of a dissociation between spatial and temporal representations in cortex of regions about the blind spot. It was long speculated that spatial completions across a blind spot were explicable simply by there being abutting representations in cortex, with no representation of the intervening space [20, 21]. Studies of spatial perception, however, have shown that representations that complete across a physiological blind spot almost *entirely* encapsulate the extent of blindness [6–9], establishing that spatial vision can generate representations that encapsulate the extent of blindness associated with the blind spot. Our data suggest an absence of a similar allowance in temporal perception.

We deliberately adopted presentation protocols, in both experiments, with simultaneous stimulations on either side of a physiological blind spot. In studies of spatial vision we would argue that such stimulation has been essential to revealing situations wherein representations generated by the brain *obviously* encapsulate the extent of blindness, as opposed to merely having a representation that appears joined due to an absence of sampling *within* the blind spot [20, 21]. We have, however, found no evidence for a temporal analogue of such representations. As such, to some extent our data could be said to stand in conflict with the results of one study that examined motion-induced modulations of visibility about the blind spot [10]. That study was said to have demonstrated that when a moving pattern extended *into* the blind spot, a representation of that stimulus was generated that was extrapolated *into* the blind spot, and hence did not rely on retinal sampling. That would seem to suggest a process which generates temporal representations *within* the blind spot – and this suggestion is at least *qualitatively* at odds with our findings. We believe there is a more parsimonious interpretation of data from that paper.

Experiments seeking to examine vision about physiological blind spots face the difficulties of both eye movements, which will slightly shift the associated region of blindness from trial to trial [22], and the fact that the drop off in photosensitivity is not a step function at the blind spot edges, but rather degrades across a short distance of retina [12]. Consequently, any estimate of the edges of a physiological blind spot should be regarded as approximate (including ours). How then should data suggesting that a representation can be seen further toward the centre of a blind spot in one condition be regarded [10]? One possible interpretation is that the baseline condition has accurately detected the absolute limit of photosensitivity at the retina, and that any extension past that point must be a consequence of a process specific to the other test condition (which does not rely on sampling by the retinal mosaic). Another interpretation would be that the other condition better taps the limited photosensitivity immediately abutting the blind spot than did the baseline task [13]. We believe the second interpretation should be preferred in the absence of additional evidence (particularly if the apparent extension is slight), as it does not assume visual representations that do not rely on sampling by the retinal mosaic.

We sought to avoid such difficulties by examining interactions involving inputs that were clearly visible to either side of a blind spot – a situation that seems somewhat more analogous to studies of spatial vision about the blind spot [20, 21]. A major difference in Experiment 1, however, was that we deliberately chose a stimulus configuration that we felt would be unlikely to complete into a perceptually bound representation *across* the blind spot – with different coloured and sized elements to either side of the blind spot (see Figure 1a). Subjectively, this precaution seemed successful, with perceptual experiences during pilot testing consisting of two clearly and persistently separate elements that seemed to collide on most trials (except trials when the initially static element began moving *before* the ‘collision’). This anecdotal evidence is supported by the fact that participants in the formal experiment did not report more element deformations or speed changes for blind spot tests relative to control site tests. This leaves open the possibility of additional processes that might only be triggered when elements *do* complete across the blind spot. From this perspective it would be interesting to conduct further investigations of the pre-conditions for elements seeming to complete *across* the blind spot, and to examine how these circumstances might impact not only spatial, but temporal vision.

In Experiment 2 we used identically coloured and shaped elements on either side of the blind spot, and subjectively these seemed to complete across the blind spot when they were flashed in tandem. We would therefore suggest that we have examined a situation where perceptual completion across the blind spot *was* triggered. Regardless, we found evidence for a weakening of brain activity entrained by flashed elements at the blind spot, relative to a control site matched in terms of eccentricity and visual hemi-field. This suggests that dynamics on either side of a physiological blind spot are encoded by a smaller pool of neurons, relative to the control site where physical stimulation was equally separated on the retina. These data are therefore consistent with the dynamics on either side of a physiological blind spot being encoded by a pool of neurons that have at least partially overlapping receptive fields. We think it is reasonable to speculate that spatial vision could be similarly informed by neurons with receptive fields that encompass either side of a physiological blind spot, and that these could be involved in triggering perceptual completions across the blind spot. Although we hasten to add that in spatial vision, any such triggered representation seems to encapsulate the majority of the spatial extent of blindness, in a manner that does not seem to be reflected in temporal vision in our experiments.

The issues we are examining touch on a broader debate in perception – to what extent are sensory experiences extrapolated from input? As Helmholtz [23] anticipated, our sensory experiences are not just governed by input, but by expectations built up recently, across our lifetimes and, potentially, our evolutionary history. These do not just shape interpretations of input [24], but can govern sensitivity [25]. Such findings have motivated claims that human vision creates perceptually explicit representations that *extrapolate* inputs into predicted locations based on preceding input dynamics [26], including extrapolations into the blind spot [10]. We are sceptical of such suggestions, as in the real-world movement can contradict expectations, with sudden stoppages and trajectory changes. If perceptual experiences were extrapolated, these would seemingly have to be erased from memory when those predictions prove to be erroneous [27–29]. This additional sensory process – of perceptual extrapolation – also seems unnecessary as it is *certain* that motor planning *does* compute predicted trajectories, otherwise we would never be able to intercept or avoid a moving object [30]. We therefore prefer more parsimonious interpretations of data that do not assume perceptually explicit extrapolations of perceived location based on movement, unless this is necessary to understand data. We are not aware of any circumstance where this has proven necessary. All instances where a perceptual extrapolation based on movement has been postulated [26, 31] seem to be equally or more explicable via explanatory frameworks that do not assume a perceptual extrapolation [27, 29, 32].

While our data touch on broader issues in sensory neuroscience, our specific interest is the possibility of there being a process of temporal filling-in at the blind spot, analogous to the filling-in observed in spatial vision – which can encapsulate the extent of blindness [20, 21]. We find no such evidence. Instead, our data are consistent with the dynamics of input on either side of a physiological blind spot being encoded by a pool of neurons with overlapping receptive fields. In sum, on the basis of our data, we would argue that we have evidence for an absence of temporal filling-in at the physiological blind spot.

## Supporting information

Experimental supplementary code

## ACKNOWLEDGEMENTS

This work was supported by an Australian Research Council Discovery Grant awarded to DHA.

## AUTHOR CONTRIBUTIONS

D.H.A. and N.D. programmed the experiments and analysed data. N.D. tested participants and wrote the initial draft of the manuscript. D.H.A. and D.S.S. edited the later versions of manuscripts. All authors reviewed the final manuscript.

## COMPETING INTERESTS

The authors declare no competing interests.

## DATA AVAILABILITY

The datasets generated and analysed during the current study will be made available via UQeSpace.

## Notes

### Competing Interest Statement

The authors have declared no competing interest.

