## Supplementary material for "An absence of temporal filling-in at the physiological blind-spot": Experimental supplementary code

Running Title: An absence of temporal filling-in at the physiological blind-spot

Submission Date: October 2020

Corresponding Author: Neha Dhupia

**Experimental codes:**

**Experiment 1: Causality Perception**

```
close all
clear all

commandwindow

sName = 'Test';   % Enter participant's initials


%% Open output file...
fid =
fopen('/Users/psychology/Desktop/Cloudstor/Blindspot_Filling_In/Chapter
1/Blindspot_Filling_In_Causality_Perception/Blindspot_Filling_In_Output
/Blindspot_Filling_In_VisLab.txt', 'a');

%fid = fopen('/Users/psychology/Dropbox/VisLab_Surface1/Shared/UQ
Perception Lab/Blindspot_Filling_In/Blindspot_Filling_In.txt', 'a');

fprintf(fid, '\n\n\n\nParticipant: %s', sName); %Insert A new line and
print subject's name

Time = datestr(now);
fprintf(fid, '\n\nStarted Session: %s \n', Time);



PsychDefaultSetup(2);
%If there are multiple displays guess that one without the menu bar is
% Setup Psychtoolbox the

%best choice.  Dislay 0 has the menu bar.

HideCursor;

screens=Screen('Screens');
screenNumber=max(screens);

Screen('Preference', 'SkipSyncTests', 1);
%PsychDebugWindowConfiguration

%Open a window.  Note the new argument to OpenWindow with value 2,
%specifying the number of buffers to the onscreen window.
[window,windowRect]=Screen(screenNumber,'OpenWindow', 0,[],[],2);


%Compute display dimensions:
[width,height] = RectSize(windowRect);


%Give the display a moment to recover from the change of display mode
when
%opening a window. It takes some monitors and LCD scan converters a few
seconds to resync.
WaitSecs(2);
```

```
scrnWidthPix=windowRect(3)-windowRect(1); % width of screen in pixels
scrnHeightPix=windowRect(4)-windowRect(2); % width of screen in pixels
scrnCenterX=windowRect(1)+(windowRect(3)-windowRect(1))/2 % horizontal
center of screen in pixels
scrnCenterY=windowRect(2)+(windowRect(4)-windowRect(2))/2 % vertical
center of screen in pixels

Screen_Width_CMs = 36;
Screen_Height_CMs = 26.5;

Viewing_Distance_CMs = 57;

Pixels_per_DVA = round((scrnWidthPix/Screen_Width_CMs) *
(Viewing_Distance_CMs/57));
Pixels_per_DVA_height = round((scrnHeightPix/Screen_Height_CMs) *
(Viewing_Distance_CMs/57));

X1=100;
Y1=height/3;

dotColor = [1 1 1];
dot1SizePix = 20; %100;

Bar_Width = 12*Pixels_per_DVA;
Bar_Height = 0.5*Pixels_per_DVA;

%% Instructions...
Screen('TextFont',window, 'Courier New');
Screen('TextSize',window, 40);
Screen('TextStyle', window, 1+2);
Screen('DrawText', window, 'Instructions...', 0, 75, [255, 255, 255,
255]);

%Draw Text
Screen('TextSize',window, 16);
Screen('DrawText', window, 'This quick procedure (4 trials) will locate
the approximate position of your blindspot.', 50, 150, [255, 255, 255,
255]);
Screen('DrawText', window, 'To begin each trial, you have to slide a
cursor into a small ring, by pressing the LEFT mouse', 50, 200, [255,
255, 255, 255]);
Screen('DrawText', window, 'button and moving the mouse to slide the
cursor.', 50, 250, [255, 255, 255, 255]);

Screen('DrawText', window, 'Once the cursor enters the ring, it will
turn green and stop moving. Please let go of the mouse button', 50,
300, [255, 255, 255, 255]);
Screen('DrawText', window, 'and keep looking at the green cursor
throughtout the trial!', 50, 350, [255, 255, 255, 255]);

Screen('DrawText', window, 'When the cursor turns green, a moving disc
will appear. DONT LOOK AT IT!', 50, 425, [255, 255, 255, 255]);
Screen('DrawText', window, 'Keep fixating the static green cursor, and
press the LEFT mouse button again as soon as the moving disc', 50, 500,
[255, 255, 255, 255]);
```

```
Screen('DrawText', window, 'dissapears. Reaction time is important, so
please press the mouse button as soon as the disc dissapears,', 50,
550, [255, 255, 255, 255]);
Screen('DrawText', window, 'but NOT before!', 50, 1200, [255, 255, 255,
255]);

Screen('DrawText', window, 'Before you start, please make sure that you
are wearing an eye patch over your LEFT eye.', 50, 750, [255, 255, 255,
255]);

Screen('DrawText', window, 'When you are ready to start this procedure,
click the LEFT mouse button...', 50, 800, [255, 255, 255, 255]);

Screen(window, 'Flip');

buttons = 0; % When the user clicks the mouse, 'buttons' becomes
nonzero.
while buttons == 0
    [mX, mY, buttons] = GetMouse; % Check for a response...
    %T_Position = randperm(3,1);
end;


% Flip to the screen
Screen('Flip', window);


%% Initial blindspot localiser routine...
BS_XYs = [];

% Set Dot Translation Speed
Dot_Speed_per_sec = width/15;

% Loop the animation until a key is pressed
%while ~KbCheck
[mX,mY,buttons]=GetMouse;
while sum(buttons)~= 0
    [mX,mY,buttons]=GetMouse;
end

Preliminary_BS_Coords = zeros(1,4);
First_Guess_X = 0;
for BS_Est = 1 : 12

    BS_Est_Num = 1+mod(BS_Est-1,4);

    Repeat = 0;

    while Repeat == 0


        %% Sequence to initiate trial...
        SetMouse(width/2,height/2);
        [mX,mY,buttons]=GetMouse;
        while sum(buttons)~= 0
            [mX,mY,buttons]=GetMouse;
        end
        stop = 0;
```

```
        StartX=0;
        StartY=0;
        NowY=0;
        NowX=0;

        count=0;

        rand_pos1 = rand(1);
        if rand_pos1 < 0.5
            Y2 = Y1-Pixels_per_DVA;
        else
            Y2 = Y1+Pixels_per_DVA;
        end
        pause(0.1)

        while stop == 0
            [mX,mY,buttons]=GetMouse;

            if  Y1-(Y2-(StartY-NowY)) < -6 ||  Y1-(Y2-(StartY-NowY)) >
6
                Colour = [255, 0, 0, 0];
            else
                Colour = [0, 255, 0, 0];
                stop=1;
            end

            if sum(buttons) > 0 && count == 0
                StartX = mX;
                StartY = mY;
                NowX = mX;
                NowY=mY;
                count=count+1;
            elseif sum(buttons) > 0
                NowX=mX;
                NowY=mY;
            end
            Screen('DrawDots', window, [X1-(StartX-NowX) Y2-(StartY-
NowY)] , dot1SizePix, Colour, [], 2);
            Screen('FrameOval', window, Colour, [X1-Pixels_per_DVA/4
Y1-Pixels_per_DVA/4 X1+Pixels_per_DVA/4 Y1+Pixels_per_DVA/4], 3, 3);
            Screen('Flip', window);
        end
        Screen('FillOval', window, Colour, [100-Pixels_per_DVA/4
(height/3)-Pixels_per_DVA/4 100+Pixels_per_DVA/4
(height/3)+Pixels_per_DVA/4]);
        Screen('Flip', window);
        [mX,mY,buttons]=GetMouse;
        while sum(buttons)~= 0
            [mX,mY,buttons]=GetMouse;
        end


        %% Preliminary BS coordinate estimate procedure
        time = 0;
        tstrat = tic;
        while sum(buttons)== 0

            switch BS_Est_Num
```

```
                case 1
                    % Testing RIGHT eye
                    Xpos = X1 + (Dot_Speed_per_sec*time);
                    Ypos = Y1+100;
                case 2
                    % Testing RIGHT eye
                    Xpos = scrnWidthPix-dot1SizePix -
(Dot_Speed_per_sec*time);
                    Ypos = Y1+100;
                case 3
                    Xpos = First_Guess_X/2;
                    Ypos = dot1SizePix + (Dot_Speed_per_sec*time);
                case 4
                    Xpos = First_Guess_X/2;
                    Ypos = (height*0.75-dot1SizePix) -
(Dot_Speed_per_sec*time);

            end

            Screen('FillOval', window, Colour, [100-Pixels_per_DVA/4
(height/3)-Pixels_per_DVA/4 100+Pixels_per_DVA/4
(height/3)+Pixels_per_DVA/4]);
            Screen('FillOval', window, [0, 255, 0, 0], [Xpos-
dot1SizePix/2, Ypos-dot1SizePix/2, Xpos+dot1SizePix/2,
Ypos+dot1SizePix/2] );

            % Flip to the screen
            Screen(window, 'Flip');

            % Increment the time
            time = toc(tstrat);

            [mX,mY,buttons]=GetMouse;

        end

        while sum(buttons)>0
            [mX,mY,buttons]=GetMouse;
        end

        %% Check Response...
        Screen('TextFont',window, 'Courier New');
        Screen('TextStyle', window, 1+2);
        Screen('TextSize',window, 16);
        Screen('DrawText', window, 'Did you click as soon as the disc
dissapeared, but not before (Left Click)', 50, Y1-25, [255, 255, 255,
255]);
        Screen('DrawText', window, 'Or did you click early or late
(Right Click)', 50, Y1+25, [255, 255, 255, 255]);
        % Flip to the screen
        Screen('Flip', window);

        % buttons = [0 0 0]
        while sum(buttons)<1
            [mX,mY,buttons]=GetMouse;
        end
        if buttons(1) == 1
            Repeat = 1;
```

```
        end
        Screen('Flip', window);


    end

    if BS_Est <= 2
        First_Guess_X = First_Guess_X+Xpos;
    elseif BS_Est > 4
        if BS_Est_Num < 3
            Preliminary_BS_Coords(BS_Est_Num) =
Preliminary_BS_Coords(BS_Est_Num)+Xpos;
        else
            Preliminary_BS_Coords(BS_Est_Num) =
Preliminary_BS_Coords(BS_Est_Num)+Ypos;
        end
    end

end

% X and Y coordinates for Left'
BS_XYs(1,1) = Preliminary_BS_Coords(1)/2;

% X and Y coordinates for Right'
BS_XYs(2,1) = Preliminary_BS_Coords(2)/2;

% X and Y coordinates for Top'
BS_XYs(1,2) = Preliminary_BS_Coords(3)/2;

% X and Y coordinates for Bottom'
BS_XYs(2,2) = Preliminary_BS_Coords(4)/2;


BS_Centre = [BS_XYs(1,1)+((BS_XYs(2,1)-BS_XYs(1,1))/2)
BS_XYs(1,2)+((BS_XYs(2,2)-BS_XYs(1,2))/2)] ;
BS_Width = BS_XYs(2,1)-BS_XYs(1,1);
BS_Height = BS_XYs(2,2)-BS_XYs(1,2);


while sum(buttons)>0
    [mX,mY,buttons]=GetMouse;
end

%% Instructions...
Screen('TextFont',window, 'Courier New');
Screen('TextSize',window, 40);
Screen('TextStyle', window, 1+2);
Screen('DrawText', window, 'Instructions...', 0, 75, [255, 255, 255,
255]);

%Draw Text
Screen('TextSize',window, 16);
Screen('DrawText', window, 'In the next quick procedure you need to
report when moving discs APPEAR.', 50, 200, [255, 255, 255, 255]);
Screen('DrawText', window, 'Again, to begin each trial, you have to
slide a cursor into a small ring.', 50, 250, [255, 255, 255, 255]);
```

```
Screen('DrawText', window, 'Some time after the cursor turns green, a moving disc will appear.', 50, 350, [255, 255, 255, 255]);
Screen('DrawText', window, 'Keep fixating the static green cursor, and press the LEFT mouse button as soon as the moving disc', 50, 400, [255, 255, 255, 255]);
Screen('DrawText', window, 'appears. Reaction time is important, so please press the mouse button as soon as the disc appears,', 50, 450, [255, 255, 255, 255]);
Screen('DrawText', window, 'but NOT before!', 50, 500, [255, 255, 255, 255]);

Screen('DrawText', window, 'When you are ready to start this procedure, click the LEFT mouse button...', 50, 660, [255, 255, 255, 255]);

Screen(window, 'Flip');

while sum(buttons)<1
    [mX,mY,buttons]=GetMouse;
end


% Flip to the screen
Screen('Flip', window);


%% Initial blindspot localiser routine %% inside out moving dot...
BS_XYs2 = [];


% Loop the animation until a key is pressed
%while ~KbCheck
[mX,mY,buttons]=GetMouse;
while sum(buttons)~= 0
    [mX,mY,buttons]=GetMouse;
end

Preliminary_BS_Coords=[];
for BS_Est = 1 : 8

    BS_Est_Num = 1+mod(BS_Est-1,4);

    Repeat = 0;
    while Repeat == 0


        if BS_Est_Num == 1
            Ypos = Y1+100; %yCenter; %yaha change kiya
        elseif BS_Est_Num == 2
            % Testing RIGHT eye
            tX = scrnWidthPix-dot1SizePix;
        elseif BS_Est_Num == 3
            'X and Y coordinates for Top'
            BS_XYs(1,2) = Ypos;
        else
            'X and Y coordinates for Bottom'
            BS_XYs(2,2) = Ypos;
        end
```

```
        %% Sequence to initiate trial...
        SetMouse(width/2,height/2);
        [mX,mY,buttons]=GetMouse;
        while sum(buttons)~= 0
            [mX,mY,buttons]=GetMouse;
        end
        stop = 0;
        StartX=0;
        StartY=0;
        NowY=0;
        NowX=0;

        count=0;

        rand_pos1 = rand(1);
        if rand_pos1 < 0.5
            Y2 = Y1-Pixels_per_DVA; %yaha change kiya hai
        else
            Y2 = Y1+Pixels_per_DVA; % yaha change kiya hai
        end
        pause(0.1)

        while stop == 0
            [mX,mY,buttons]=GetMouse;

            if  Y1-(Y2-(StartY-NowY)) < -6 ||  Y1-(Y2-(StartY-NowY)) >
6
                Colour = [255, 0, 0, 0];
            else
                Colour = [0, 255, 0, 0];
                stop=1;
            end

            if sum(buttons) > 0 && count == 0
                StartX = mX;
                StartY = mY;
                NowX = mX;
                NowY=mY;
                count=count+1;
            elseif sum(buttons) > 0
                NowX=mX;
                NowY=mY;
            end
            Screen('DrawDots', window, [X1-(StartX-NowX) Y2-(StartY-
NowY)] , dot1SizePix, Colour, [], 2);
            Screen('FrameOval', window, Colour, [X1-Pixels_per_DVA/4
Y1-Pixels_per_DVA/4 X1+Pixels_per_DVA/4 Y1+Pixels_per_DVA/4], 3, 3);
            Screen('Flip', window);
        end
        Screen('FillOval', window, Colour, [100-Pixels_per_DVA/4
(height/3)-Pixels_per_DVA/4 100+Pixels_per_DVA/4
(height/3)+Pixels_per_DVA/4]);
        Screen('Flip', window);
        [mX,mY,buttons]=GetMouse;
        while sum(buttons)~= 0
            [mX,mY,buttons]=GetMouse;
        end
```

```
        %% Preliminary BS coordinate estimate procedure
        X2 = BS_Centre(1);
        Y2 = BS_Centre(2);
        tX2 = X2-dot1SizePix;


        time = 0;
        tstrat = tic;
        while sum(buttons)== 0

            switch BS_Est_Num
                case 1
                    % Testing RIGHT eye
                    Xpos2 = X2 + (Dot_Speed_per_sec*time);
                    Ypos = BS_Centre(2);

                case 2
                    % Testing RIGHT eye
                    Xpos2 = tX2 - (Dot_Speed_per_sec*time);
                    Ypos = BS_Centre(2);

                case 3
                    Xpos2 = BS_Centre(1);
                    Ypos = (Y2-dot1SizePix) + (Dot_Speed_per_sec*time);

                case 4
                    Xpos2 = BS_Centre(1);
                    Ypos = (Y2-dot1SizePix) - (Dot_Speed_per_sec*time);

            end


            Screen('FillOval', window, Colour, [100-Pixels_per_DVA/4
(height/3)-Pixels_per_DVA/4 100+Pixels_per_DVA/4
(height/3)+Pixels_per_DVA/4]);
            Screen('DrawDots', window, [X2 Y2] , dot1SizePix, [255 255
255], [], 2);
            Screen('FillOval', window, [0, 255, 0, 0], [Xpos2-
dot1SizePix/2, Ypos-dot1SizePix/2, Xpos2+dot1SizePix/2,
Ypos+dot1SizePix/2] );

            % Flip to the screen
            Screen(window, 'Flip');

            % Increment the time
            time = toc(tstrat);

            [mX,mY,buttons]=GetMouse;

        end

        while sum(buttons)>0
            [mX,mY,buttons]=GetMouse;
        end

        %% Check Response...
```

```
        Screen('TextFont',window, 'Courier New');
        Screen('TextStyle', window, 1+2);
        Screen('TextSize',window, 16);
        Screen('DrawText', window, 'Did you click as soon as the disc
appeared, but not before (Left Click)', 50, Y1-25, [255, 255, 255,
255]);
        Screen('DrawText', window, 'Or did you click early or late
(Right Click)', 50, Y1+25, [255, 255, 255, 255]);
        % Flip to the screen
        Screen('Flip', window);

        % buttons = [0 0 0]
        while sum(buttons)<1
            [mX,mY,buttons]=GetMouse;
        end
        if buttons(1) == 1
            Repeat = 1;
        end
        Screen('Flip', window);

    end

    if BS_Est > 4
        if BS_Est_Num == 1
            'X and Y coordinates for Left'
            BS_XYs2(1,1) = Xpos2;
        elseif BS_Est_Num == 2
            'X and Y coordinates for Right'
            BS_XYs2(2,1) = Xpos2;
        elseif BS_Est_Num == 3
            'X and Y coordinates for Top'
            BS_XYs2(1,2) = Ypos;
        else
            'X and Y coordinates for Bottom'
            BS_XYs2(2,2) = Ypos;
        end
    end

end


BS_XYs_Avg(1,1)=(BS_XYs2(1,1)+BS_XYs(1,1))/2;  %Right
BS_XYs_Avg(2,1)=(BS_XYs2(2,1)+BS_XYs(2,1))/2;  %Left
BS_XYs_Avg(1,2)=(BS_XYs2(1,2)+BS_XYs(1,2))/2;  %Bottom
BS_XYs_Avg(2,2)=(BS_XYs2(2,2)+BS_XYs(2,2))/2;  %Top

BS_Centre2 = [BS_XYs_Avg(1,1)+((BS_XYs_Avg(2,1)-BS_XYs_Avg(1,1))/2)
BS_XYs_Avg(1,2)+((BS_XYs_Avg(2,2)-BS_XYs_Avg(1,2))/2)] ;
BS_Width2 = BS_XYs_Avg(1,1)-BS_XYs_Avg(2,1);
BS_Height2 = BS_XYs_Avg(1,2)-BS_XYs_Avg(2,2);


while sum(buttons)>0
    [mX,mY,buttons]=GetMouse;
end

%% Instructions for Fill-in...
```

```
Screen('TextFont',window, 'Courier New');
Screen('TextSize',window, 40);
Screen('TextStyle', window, 1+2);
Screen('DrawText', window, 'Instructions...', 0, 75, [255, 255, 255,
255]);

%Draw Text
Screen('TextSize',window, 16);
Screen('DrawText', window, 'This procedure will measure the position of
your physiological blindspot more precisely.', 50, 250, [255, 255, 255,
255]);
Screen('DrawText', window, 'To begin each trial, again you have to
slide a cursor into a small ring, and it will turn green.', 50, 350,
[255, 255, 255, 255]);

Screen('DrawText', window, 'Once the cursor turns green, keep looking
at it.', 50, 450, [255, 255, 255, 255]);
Screen('DrawText', window, 'Then there will be two green bars. At first
these will be clearly seperated. Your task is make them', 50, 500,
[255, 255, 255, 255]);
Screen('DrawText', window, 'stretch (by pressing the LEFT mouse button)
until they seem to touch. As soon as they TOUCH, release', 50, 550,
[255, 255, 255, 255]);
Screen('DrawText', window, 'the LEFT mouse button immediately (timing
is important). Then you should press the MIDDLE mouse', 50, 600, [255,
255, 255, 255]);
Screen('DrawText', window, 'button, and the next trial will begin...',
50, 650, [255, 255, 255, 255]);


Screen('DrawText', window, 'When you are ready to start this procedure,
click the LEFT mouse button...', 50, 750, [255, 255, 255, 255]);

Screen(window, 'Flip');

while sum(buttons)<1
    [mX,mY,buttons]=GetMouse;
end


% Flip to the screen
Screen('Flip', window);

%% Initial fill-in ...


Element_Wd = 2;
Element_Hgt = 0.5;

Animation_Duration = 1;
Start_Distance = 3;
Travel_Distance = 5;

Num_Trials = 10;


%%
BS_Coords = BS_Centre2;
```

```
BS_Coords(3) = BS_Width2;
BS_Coords(4) = BS_Height2;

% Save_Struct{1}=BS_Coords(1);
% Save_Struct{2}=BS_Coords(2);
% Save_Struct{3}=BS_Coords(3);
% Save_Struct{4}=BS_Coords(4);
%
% savefile = 'BS_Coords.mat';
% save(savefile, 'Save_Struct');


%% Load BS coordinates

% FileName = 'BS_Coords.mat';
% BS_Coords = load(FileName);
% BS_Coords = BS_Coords.Save_Struct;


for count = 1:Num_Trials


    %% Sequence to initiate trial...
    SetMouse(width/2,height/2);
    [mX,mY,buttons]=GetMouse;
    while sum(buttons)~= 0
        [mX,mY,buttons]=GetMouse;
    end
    stop = 0;
    StartX=0;
    StartY=0;
    NowY=0;
    NowX=0;

    init_count=0;

    rand_pos1 = rand(1);
    if rand_pos1 < 0.5
        Y2 = Y1-Pixels_per_DVA;
    else
        Y2 = Y1+Pixels_per_DVA;
    end
    pause(0.1)

    %[Start_MouseX,Start_MouseY,buttons]=GetMouse;
    while stop == 0
        [mX,mY,buttons]=GetMouse;

        if  Y1-(Y2-(StartY-NowY)) < -6 ||  Y1-(Y2-(StartY-NowY)) > 6
            Colour = [255, 0, 0, 0];
        else
            Colour = [0, 255, 0, 0];
            stop=1;
        end

        if sum(buttons) > 0 && init_count == 0
            StartX = mX;
            StartY = mY;
```

```
            NowX = mX;
            NowY=mY;
            init_count=init_count+1;
        elseif sum(buttons) > 0
            NowX=mX;
            NowY=mY;
        end
        %NowX = Start_MouseX-mX;
        Screen('DrawDots', window, [X1-(StartX-NowX) Y2-(StartY-NowY)]
, dot1SizePix, Colour, [], 2);
        %Screen('DrawDots', windowPtr, xy [,size] [,color] [,center]
[,dot_type][,lenient]);
        Screen('FrameOval', window, Colour, [X1-Pixels_per_DVA/4 Y1-
Pixels_per_DVA/4 X1+Pixels_per_DVA/4 Y1+Pixels_per_DVA/4], 3, 3);
        Screen('Flip', window);
    end
        Screen('FillOval', window, Colour, [100-Pixels_per_DVA/4
(height/3)-Pixels_per_DVA/4 100+Pixels_per_DVA/4
(height/3)+Pixels_per_DVA/4]);
        Screen('Flip', window);


 %% Test protocol...
    BS_Colour = [255, 0, 0, 0];
    Screen('FillOval', window, Colour, [100-Pixels_per_DVA/4
(height/3)-Pixels_per_DVA/4 100+Pixels_per_DVA/4
(height/3)+Pixels_per_DVA/4]);

    SetMouse(width/2,height/2);
    [mX,mY,buttons]=GetMouse;
    while sum(buttons)~= 0
        [mX,mY,buttons]=GetMouse;
    end

    stop = 0;



    BS_Colour = [255, 0, 0, 0];
    Stop = 0;
    Wd = 11.75;
    while Stop == 0
        [mX,mY,buttons]=GetMouse;
        Screen('FillOval', window, Colour, [100-Pixels_per_DVA/4
(height/3)-Pixels_per_DVA/4 100+Pixels_per_DVA/4
(height/3)+Pixels_per_DVA/4]);
        Screen('FillRect', window, [0 255 0], [BS_Coords(1)-
(6*Pixels_per_DVA) BS_Coords(2)-(Element_Hgt/2)*Pixels_per_DVA
BS_Coords(1)+(6*Pixels_per_DVA)
BS_Coords(2)+(Element_Hgt/2)*Pixels_per_DVA]);
        Screen('FillRect', window, [0 0 0], [BS_Coords(1)-
(Wd/2*Pixels_per_DVA) BS_Coords(2)-(Element_Hgt/2)*Pixels_per_DVA
BS_Coords(1)+(Wd/2*Pixels_per_DVA)
BS_Coords(2)+(Element_Hgt/2)*Pixels_per_DVA]);

        if buttons(3) == 1
            Stop = 1;
        elseif buttons(1) == 1
```

```
            Wd = Wd-0.05;
        elseif buttons(2) == 1
            Wd = Wd+0.05;
        end

        if Wd > 11.75
            Wd = 11.75;
        elseif Wd < 1
            Wd = 1;
        end
        Screen('Flip', window);
    end


    [mX,mY,buttons]=GetMouse;
    while sum(buttons)~= 0
        [mX,mY,buttons]=GetMouse;
    end

    Results_Array(count) = Wd;

    fprintf(fid, '\nWidth Setting %5.2f', Wd);

end

Mean_Wd = mean(Results_Array);


%% Instructions Temporal filling In...

Screen('TextFont',window, 'Courier New');
Screen('TextSize',window, 40);
Screen('TextStyle', window, 1+2);
Screen('DrawText', window, 'Instructions...', 0, 75, [255, 255, 255,
255]);

%Draw Text
Screen('TextSize',window, 16);
Screen('DrawText', window, 'Now we are going to take a TIMING
measure.', 50, 150, [255, 255, 255, 255]);

Screen('DrawText', window, 'Again, to begin each trial you have to
slide the cursor into the ring.', 50, 200, [255, 255, 255, 255]);

Screen('DrawText', window, 'When the cursor turns green, a green moving
and a larger red static bar will', 50, 250, [255, 255, 255, 255]);
Screen('DrawText', window, 'appear. DONT LOOK AT THEM! Keep fixating
the static green cursor.', 50, 300, [255, 255, 255, 255]);

Screen('DrawText', window, 'After some time, the green bar will stop,
and the larger red bar will begin moving.', 50, 400, [255, 255, 255,
255]);
Screen('DrawText', window, 'Your task is to say if the red bar seemed
to get HIT, and to be launched into motion,', 50, 450, [255, 255, 255,
255]);
Screen('DrawText', window, 'by the green bar, like a pool ball being
hit by a cue.', 50, 500, [255, 255, 255, 255]);
```

```
Screen('DrawText', window, 'Alternatively, the red bar may seem to
start moving without being HIT,', 50, 550, [255, 255, 255, 255]);
Screen('DrawText', window, 'or it may seem to start moving AFTER it
gets HIT by the green bar.', 50, 600, [255, 255, 255, 255]);

Screen('DrawText', window, 'When you are ready to begin, click the LEFT
mouse button...', 50, 700, [255, 255, 255, 255]);

Screen(window, 'Flip');

buttons = 0; % When the user clicks the mouse, 'buttons' becomes
nonzero.
while buttons == 0
    [mX, mY, buttons] = GetMouse; % Check for a response...
    %T_Position = randperm(3,1);
end

% Flip to the screen
Screen('Flip', window);


%% Initial Temporal filling-in ...


Adjacent = BS_Coords(1)-100;
Opposite = BS_Coords(2)-(height/3);
Hypotenuse = sqrt((Adjacent*Adjacent)+(Opposite*Opposite));
Angle = atand(Opposite/Adjacent);

Control_Angle = Angle-60;

Control_X = cosd(Control_Angle)*Hypotenuse;
Control_Y = -(sind(Control_Angle)*Hypotenuse);

Check_Angle = atand(Control_X/Control_Y)-Control_Angle
Check_Hypotenuse = sqrt((Control_Y*Control_Y)+(Control_X*Control_X))-
Hypotenuse




Time = datestr(now);
fprintf(fid, '\n\nStarted Main Experiment: %s \n', Time);

ISIs = -1:0.25:1;
Temporal_Results = zeros(size(ISIs,2),3,3,2);
Temporal_Results(:,1,1,1)=ISIs;
Temporal_Results(:,1,2,1)=ISIs;
Temporal_Results(:,1,3,1)=ISIs;
Temporal_Results(:,1,1,2)=ISIs;
Temporal_Results(:,1,2,2)=ISIs;
Temporal_Results(:,1,3,2)=ISIs;

N_Repeats = 9;

Num_Trials = size(ISIs,2)*N_Repeats*2;

Change_Report = [0 0];
```

```
% Get framerate
Hz = FrameRate(window);

Trial_Order = randperm(Num_Trials);

Animation_Duration = 4;
for count = 1:Num_Trials

    Test_Number = 1+mod(Trial_Order(count),size(ISIs,2))
    Test_ISI = ISIs(Test_Number)

    if Trial_Order(count)<= Num_Trials/2
        % Testing at blindspot
        Condition = 1
        Test_Y = BS_Coords(2);
%         BS_Rt_Side =
BS_Coords(1)+(BS_Coords(3)/2)+(4*Pixels_per_DVA)-
(Mean_Wd*Pixels_per_DVA);
%         BS_Rt_Side =
BS_Coords(1)+(BS_Coords(3)/2)+(4*Pixels_per_DVA)-
(Mean_Wd*Pixels_per_DVA);
        BS_Lt_Side = BS_Coords(1)-(Mean_Wd/2*Pixels_per_DVA);
        BS_Rt_Side = BS_Coords(1)+(Mean_Wd/2*Pixels_per_DVA);
    else
        % Testing at control position
        Condition = 2
        Test_Y = (height/3)+Control_Y;
        BS_Rt_Side = Control_X+100;
        BS_Lt_Side = Control_X+100;
    end

    %% Sequence to initiate trial for Temporal filling In...

    SetMouse(width/2,height/2);
    [mX,mY,buttons]=GetMouse;
    while sum(buttons)~= 0
        [mX,mY,buttons]=GetMouse;
    end
    stop = 0;
    StartX=0;
    StartY=0;
    NowY=0;
    NowX=0;

    init_count=0;

    rand_pos1 = rand(1);
    if rand_pos1 < 0.5
        Y2 = (height/3)-Pixels_per_DVA;
    else
        Y2 = (height/3)+Pixels_per_DVA;
    end
    pause(0.1)

    while stop == 0
        [mX,mY,buttons]=GetMouse;
```

```
        if  (height/3)-(Y2-(StartY-NowY)) < -6 ||  (height/3)-(Y2-
(StartY-NowY)) > 6
            Colour = [255, 0, 0, 0];
        else
            Colour = [0, 255, 0, 0];
            stop=1;
        end

        if sum(buttons) > 0 && init_count == 0
            StartX = mX;
            StartY = mY;
            NowX = mX;
            NowY=mY;
            init_count=init_count+1;
        elseif sum(buttons) > 0
            NowX=mX;
            NowY=mY;
        end
        Screen('DrawDots', window, [100-(StartX-NowX) Y2-(StartY-NowY)]
, dot1SizePix, Colour, [], 2);
        Screen('FrameOval', window, Colour, [100-Pixels_per_DVA/4
(height/3)-Pixels_per_DVA/4 100+Pixels_per_DVA/4
(height/3)+Pixels_per_DVA/4], 3, 3);
        Screen('Flip', window);
    end
        Screen('FillOval', window, Colour, [100-Pixels_per_DVA/4
(height/3)-Pixels_per_DVA/4 100+Pixels_per_DVA/4
(height/3)+Pixels_per_DVA/4]);
        Screen('Flip', window);


    %% Test protocol...

    Element_Wd = 2;
    Element_Hgt = 0.5;
%     Wd = 5;
%     Start_Distance = 10;
%     Travel_Distance = 5;

    BS_Colour = [255, 0, 0, 0];
    Screen('FillOval', window, Colour, [100-Pixels_per_DVA/4
(height/3)-Pixels_per_DVA/4 100+Pixels_per_DVA/4
(height/3)+Pixels_per_DVA/4]);
    Mov1_Pos = -5;
    Mov2_Pos = 0;
    time = 0;
    tstrat = tic;
    while time < Animation_Duration
        Screen('FillOval', window, Colour, [100-Pixels_per_DVA/4
(height/3)-Pixels_per_DVA/4 100+Pixels_per_DVA/4
(height/3)+Pixels_per_DVA/4]);

        Screen('FillRect', window, [255 0 0],
[BS_Rt_Side+(Mov2_Pos*Pixels_per_DVA) Test_Y-
(2*Element_Hgt)*Pixels_per_DVA
BS_Rt_Side+(Element_Wd*Pixels_per_DVA)+(Mov2_Pos*Pixels_per_DVA)
Test_Y+(2*Element_Hgt)*Pixels_per_DVA]);
```

```
        Screen('FillRect', window, [0 255 0], [BS_Lt_Side-
(Element_Wd*Pixels_per_DVA)-(Mov1_Pos*Pixels_per_DVA) Test_Y-
(Element_Hgt/2)*Pixels_per_DVA BS_Lt_Side-(Mov1_Pos*Pixels_per_DVA)
Test_Y+(Element_Hgt/2)*Pixels_per_DVA]);

%         Screen('FillRect', window, [255 0 0],
[BS_Rt_Side+(Mov2_Pos*Pixels_per_DVA) Test_Y-
(2*Element_Hgt)*Pixels_per_DVA
BS_Rt_Side+(Mov2_Pos*Pixels_per_DVA)+(Element_Wd*Pixels_per_DVA)
Test_Y+(2*Element_Hgt)*Pixels_per_DVA]);
%         Screen('FillRect', window, [0 255 0], [BS_Lt_Side-
(Element_Wd*Pixels_per_DVA)-(Mov1_Pos*Pixels_per_DVA) Test_Y-
(Element_Hgt/2)*Pixels_per_DVA BS_Lt_Side-(Mov1_Pos*Pixels_per_DVA)
Test_Y+(Element_Hgt/2)*Pixels_per_DVA]);


        Screen('Flip', window);
        time = toc(tstrat);
        if time <= (Animation_Duration*0.5)
            Mov1_Pos = 5 - ((time/(Animation_Duration*0.5))*5);
        end
        if time >= (Animation_Duration*0.5)+Test_ISI
            Mov2_Pos = (((time-(Animation_Duration*0.5)-Test_ISI) /
(Animation_Duration*0.5))*5);
        end
    end
    SetMouse(width/2,height/2);
    [mX,mY,buttons]=GetMouse;
    while sum(buttons)~= 0
        [mX,mY,buttons]=GetMouse;
    end

    % Increment the time
    time = toc(tstrat);

    Screen('Flip', window);

    [mX,mY,buttons]=GetMouse;
    while sum(buttons)~= 0
        [mX,mY,buttons]=GetMouse;
    end

    %% Take Response...
    Screen('TextFont',window, 'Courier New');
    Screen('TextStyle', window, 1+2);
    Screen('TextSize',window, 16);
    Screen('DrawText', window, 'Did the red bar start moving BEFORE (or
without) being HIT by the green bar (Left Click)', 50, Y1-25, [255,
255, 255, 255]);
    Screen('DrawText', window, 'Did the red bar start moving some time
AFTER being HIT by the green bar (Right Click)', 50, Y1, [255, 255,
255, 255]);
    Screen('DrawText', window, 'Or did the red bar start moving AS SOON
as it was HIT by the green bar (MIDDLE Click)', 50, Y1+25, [255, 255,
255, 255]);
    % Flip to the screen
    Screen('Flip', window);
```

```
    while sum(buttons)== 0
        [mX,mY,buttons]=GetMouse;
    end

    if buttons(1) == 1
        % Too early for launch

Temporal_Results(Test_Number,2,1,Condition)=Temporal_Results(Test_Numbe
r,2,1,Condition)+1;
        Launch_Response = -1;
    elseif buttons(2) == 1
        % Too late for launch

Temporal_Results(Test_Number,2,3,Condition)=Temporal_Results(Test_Numbe
r,2,3,Condition)+1;
        Launch_Response =  1;
    else
        % Just right for launch

Temporal_Results(Test_Number,2,2,Condition)=Temporal_Results(Test_Numbe
r,2,2,Condition)+1;
        Launch_Response = 0;
    end

Temporal_Results(Test_Number,3,1:3,Condition)=Temporal_Results(Test_Num
ber,3,1:3,Condition)+1;

    Screen('Flip', window);

    fprintf(fid, '\nTest Condition: %5.0f \tTiming: %5.2f Launch
Response: %5.0f',Condition, Test_ISI, Launch_Response);


    if mod(count,8) == 0
        % Sequence to check if moving bar seemed to change shape of
speed
        SetMouse(width/2,height/2);
        [mX,mY,buttons]=GetMouse;
        while sum(buttons)~= 0
            [mX,mY,buttons]=GetMouse;
        end
        Screen('TextSize',window, 20);
        Screen('DrawText', window, 'Did the green bar seem to change
shape or speed before it stopped moving?', 50, Y1-25, [255, 0, 0]);
        Screen('DrawText', window, 'YES (Left Click)', 50, Y1, [255,
255, 255, 255]);
        Screen('DrawText', window, 'NOT SURE (Middle Click)', 50,
Y1+25, [255, 255, 255, 255]);
        Screen('DrawText', window, 'NO (right Click)', 50, Y1+50, [255,
255, 255, 255]);
        % Flip to the screen
        Screen('Flip', window);
        while sum(buttons)== 0
            [mX,mY,buttons]=GetMouse;
        end
        if buttons(1) == 1
            Change_Report(Condition) = Change_Report(Condition)+1;
        end
```

```
        while sum(buttons)~= 0
            [mX,mY,buttons]=GetMouse;
        end
        Screen('Flip', window);
    end

end


%% Debrief...
[mX,mY,buttons]=GetMouse;
while sum(buttons)~= 0
    [mX,mY,buttons]=GetMouse;
end

Screen('TextFont',window, 'Courier New');
Screen('TextSize',window, 40);
Screen('TextStyle', window, 1+2);
Screen('DrawText', window, 'FINISHED : ) You can relax now. Thank you
for participanting!', 0, 75, [255, 255, 255, 255]);

%Draw Text
Screen('TextSize',window, 16);
Screen('DrawText', window, 'You may not have been aware of this, but
each human eye has a "blind-spot",', 50, 150, [255, 255, 255, 255]);
Screen('DrawText', window, 'a region that has no photoreceptors - so
you cannot see any images that project to it.', 50, 200, [255, 255,
255, 255]);
Screen('DrawText', window, 'But if an image, say a bar, extends right
across your blind-spot, you wont see a "hole".', 50, 250, [255, 255,
255, 255]);
Screen('DrawText', window, 'Instead, that bar will seem to complete
across the blind-spot. This process is called "perceptual', 50, 300,
[255, 255, 255, 255]);
Screen('DrawText', window, 'filling-in".', 50, 350, [255, 255, 255,
255]);

Screen('DrawText', window, 'This experiment aims to find out if your
brain fills in time, like it fills in space.', 50, 450, [255, 255, 255,
255]);
Screen('DrawText', window, 'We had an object move to the edge of your
blind-spot, so it would look like it had reached', 50, 500, [255, 255,
255, 255]);
Screen('DrawText', window, 'and touched an initially static object on
the other side. Sometimes, it should have seemed', 50, 550, [255, 255,
255, 255]);
Screen('DrawText', window, 'like a collision, that launched the
initially static object into motion. The question is', 50, 600, [255,
255, 255, 255]);
Screen('DrawText', window, 'would this seem to happen when the
initially static object began moving as soon as the initially moving',
50, 650, [255, 255, 255, 255]);
Screen('DrawText', window, 'object reached the other side of the blind-
spot, or would there need to be a delay?', 50, 700, [255, 255, 255,
255]);
```

```
Screen('DrawText', window, 'If no delay is needed, it would seem that
your brain does not fill-in time like it fills-in space', 50, 750,
[255, 255, 255, 255]);
Screen('DrawText', window, 'as it is not allowing for the time that
would be needed for the initially moving object to reach the', 50, 800,
[255, 255, 255, 255]);
Screen('DrawText', window, 'other side of your blind-spot. If a delay
is needed, it would suggest that your brain does fill-in time.', 50,
850, [255, 255, 255, 255]);
Screen('DrawText', window, 'We wont know the answer until we finish the
experiment - so thankyou for helping by participating!', 50, 900, [255,
255, 255, 255]);

Screen(window, 'Flip');




setuptestenvironment

fprintf(fid, '\n\n\n\nRESULT SUMMARIES');
fprintf(fid, '\n\nBlindspot distance from fixation: %5.1f
(dva)',(Hypotenuse / Pixels_per_DVA) );
fprintf(fid, '\n\nBlindspot X-distance from fixation: %5.1f
(dva)',(BS_Coords(1)-X1) / Pixels_per_DVA);
fprintf(fid, '\nBlindspot Y-distance from fixation: %5.1f
(dva)',(BS_Coords(2)-Y1) / Pixels_per_DVA_height);
fprintf(fid, '\nBlindspot Width: %5.1f (dva)',BS_Coords(3) /
Pixels_per_DVA);
fprintf(fid, '\nBlindspot Height: %5.1f (dva)',BS_Coords(4) /
Pixels_per_DVA_height);

for Condition = 1 : 2
    fprintf(fid, '\n\n\nTest Condition: %5.0f: \nISI \tPr. Early \tPr.
Launch \tPr. Late',Condition);
    for n = 1 : size(Temporal_Results,1)
        fprintf(fid, '\n%5.2f \t%5.2f \t\t%5.2f
\t\t%5.2f',Temporal_Results(n,1,1,Condition),
Temporal_Results(n,2,1,Condition)/Temporal_Results(n,3,1,Condition),
Temporal_Results(n,2,2,Condition)/Temporal_Results(n,3,2,Condition),
Temporal_Results(n,2,3,Condition)/Temporal_Results(n,3,3,Condition)  );
    end

    % Too Early Data
    R_Array(:,1)=Temporal_Results(:,1,1,Condition);
    R_Array(:,2)=Temporal_Results(:,2,1,Condition);
    R_Array(:,3)=Temporal_Results(:,3,1,Condition);
    R_Array(:,2) = R_Array(1,3)-R_Array(:,2);

    Early_analyses = BootstrapInference(R_Array, priors, 'nafc', 1,
'samples', 99);
    fprintf(fid, '\n\nToo Early PSE:
%5.2f',Early_analyses.thresholds(2));
    figure(1)
    GoodnessOfFit (Early_analyses)

    % Too Late Data
    R_Array(:,1)=Temporal_Results(:,1,3,Condition);
```

```
    R_Array(:,2)=Temporal_Results(:,2,3,Condition);
    R_Array(:,3)=Temporal_Results(:,3,3,Condition);
    Late_analyses = BootstrapInference(R_Array, priors, 'nafc', 1,
'samples', 99);
    fprintf(fid, '\nToo Late PSE: %5.2f',Late_analyses.thresholds(2));
    fprintf(fid, '\nCondition %4.0f PSE: %5.2f',Condition,
(Late_analyses.thresholds(2)-Early_analyses.thresholds(2))/2);
    figure(2)
    GoodnessOfFit (Late_analyses)

    fprintf(fid, '\n\nChanges reported on %5.0f percent of
trials...',(Change_Report(Condition)/(Num_Trials/2))*100);
end

Time = datestr(now);
fprintf(fid, '\n\nFinished: %s \n', Time);


status = fclose('all');


[mX,mY,buttons]=GetMouse;
while sum(buttons)== 0
    [mX,mY,buttons]=GetMouse;
end

sca
ShowCursor
```

### Experiment 2: Temporal Sensitivity

```
% close all
% clear all

commandwindow

sName = 'Test';   % Enter participant's initials

EEG_Session = 1;  % 0 = Pilot / Behavioural Testing
                    % 1 = EEG Testing session...


%% Open output file...
%fid =
fopen('/Users/psychology/Desktop/Cloudstor/Blindspot_Filling_In/Chapter
4/Output/Blindspot_Apparent_Motion_Frequency.txt', 'a');
%C:\Users\psychology\Dropbox\Research_Projects\Blindspot_Filling_In
fid =
fopen('/Users/psychology/Dropbox/Research_Projects/Blindspot_Filling_In
/Chapter
1/Blindspot_Filling_In_Apparent_Motion_EEG/Blindspot_Filling_In_AP.txt'
, 'a');
```

```
fprintf(fid, '\n\n\n\nParticipant: %s', sName); %Insert A new line and
print subject's name

Time = datestr(now);
fprintf(fid, '\n\nStarted Session: %s \n', Time);

% % % % % %% Open Visage & TDT (Just for Triggers
% % % % %
% % % % % if EEG_Session == 1
% % % % %
% % % % %     CheckCard = vsg(vsgInit,'');
% % % % %     if (CheckCard < 0)
% % % % %         return
% % % % %     end
% % % % %
% % % % %     vsg(vsgSetDrawPage,vsgVIDEOPAGE,1, 0);
% % % % %
% % % % %     RP = actxcontrol('RPco.x');
% % % % %     invoke(RP,'ConnectRP2','GB',1);
% % % % %     if invoke(RP, 'ConnectRP2','GB',1)
% % % % %          fprintf('\nTDT connected\n\n')
% % % % %     else
% % % % %          error('TDT unable to connect')
% % % % %     end
% % % % %     invoke(RP,'ClearCOF');
% % % % %
% % % % %     file =
'C:\Users\psychology\Dropbox\Research_Projects\Toolbox\Triggered_Sheppa
rd_Tone.rco';
% % % % %     check = exist(file);
% % % % %     while check == 0
% % % % %         'EMERGENCY!!!!'
% % % % %         break
% % % % %     end
% % % % %
invoke(RP,'LoadCOF','C:\Users\psychology\Dropbox\Research_Projects\Tool
box\Triggered_Sheppard_Tone.rco');
% % % % %     invoke(RP,'Run');
% % % % %
% % % % %     invoke(RP,'SetTagVal','Amp_1',0);
% % % % %     invoke(RP,'SetTagVal','Amp_2',0);
% % % % %     invoke(RP,'SetTagVal','Amp_3',0);
% % % % %     invoke(RP,'SetTagVal','Amp_4',0);
% % % % %     invoke(RP,'SetTagVal','Amp_5',0);
% % % % %
% % % % %     % TRIGGER VALS
% % % % %     Trigger_Vals = [2 8 15 50];
% % % % %
% % % % %     % Event Codes...
% % % % %     % Trigger value     Input value           Meaning
% % % % %     %
% % % % %     %             63           2 | 22        Blindspot,
Flicker Trial | Blindspot to Control, Flicker
% % % % %     %             95           8 | 88        Blindspot,
Apparent Motion Trial | Blindspot to Control, Apparent Motion Trial
% % % % %     %            127            15           Control
Position, Flicker Trial
```

```
% % % % %     %            191            50          Control
Position, Apparent Motion Trial
% % % % %
% % % % % end

%FROM HERE

PsychDefaultSetup(2);
%If there are multiple displays guess that one without the menu bar is
% Setup Psychtoolbox the
%best choice.  Dislay 0 has the menu bar.

%HideCursor;

screens=Screen('Screens');
screenNumber=max(screens);

Screen('Preference', 'SkipSyncTests', 1);
PsychDebugWindowConfiguration

%Open a window.  Note the new argument to OpenWindow with value 2,
%specifying the number of buffers to the onscreen window.
[window,windowRect]=Screen(screenNumber,'OpenWindow', 0,[],[],2);

%TO HERE


%Compute display dimensions:
[width,height] = RectSize(windowRect);


%Give the display a moment to recover from the change of display mode
when
%opening a window. It takes some monitors and LCD scan converters a few
seconds to resync.
WaitSecs(2);


scrnWidthPix=windowRect(3)-windowRect(1); % width of screen in pixels
scrnHeightPix=windowRect(4)-windowRect(2); % width of screen in pixels
scrnCenterX=windowRect(1)+(windowRect(3)-windowRect(1))/2 % horizontal
center of screen in pixels
scrnCenterY=windowRect(2)+(windowRect(4)-windowRect(2))/2 % vertical
center of screen in pixels

Screen_Width_CMs = 54;
Screen_Height_CMs = 30.5;

Viewing_Distance_CMs = 57;

Pixels_per_DVA = round((scrnWidthPix/Screen_Width_CMs) *
(Viewing_Distance_CMs/57));
Pixels_per_DVA_height = round((scrnHeightPix/Screen_Height_CMs) *
(Viewing_Distance_CMs/57));

X1=100;
Y1=height/3;
```

```
dotColor = [1 1 1];
dot1SizePix = 20; %100;

Bar_Width = 12*Pixels_per_DVA;
Bar_Height = 0.5*Pixels_per_DVA;

ShowCursor


%% Instructions...
Screen('TextFont',window, 'Courier New');
Screen('TextSize',window, 40);
Screen('TextStyle', window, 1+2);
Screen('DrawText', window, 'Instructions...', 0, 75, [255, 255, 255,
255]);

%Draw Text
Screen('TextSize',window, 16);
Screen('DrawText', window, 'This quick procedure (4 trials) will locate
the approximate position of your blindspot.', 50, 150, [255, 255, 255,
255]);
Screen('DrawText', window, 'To begin each trial, you have to slide a
cursor into a small ring, by pressing the LEFT mouse', 50, 200, [255,
255, 255, 255]);
Screen('DrawText', window, 'button and moving the mouse to slide the
cursor.', 50, 250, [255, 255, 255, 255]);

Screen('DrawText', window, 'Once the cursor enters the ring, it will
turn green and stop moving. Please let go of the mouse button', 50,
300, [255, 255, 255, 255]);
Screen('DrawText', window, 'and keep looking at the green cursor
throughtout the trial!', 50, 350, [255, 255, 255, 255]);

Screen('DrawText', window, 'When the cursor turns green, a moving disc
will appear. DONT LOOK AT IT!', 50, 425, [255, 255, 255, 255]);
Screen('DrawText', window, 'Keep fixating the static green cursor, and
press the LEFT mouse button again as soon as the moving disc', 50, 500,
[255, 255, 255, 255]);
Screen('DrawText', window, 'dissapears. Reaction time is important, so
please press the mouse button as soon as the disc dissapears,', 50,
550, [255, 255, 255, 255]);
Screen('DrawText', window, 'but NOT before!', 50, 1200, [255, 255, 255,
255]);

Screen('DrawText', window, 'Before you start, please make sure that you
are wearing an eye patch over your LEFT eye.', 50, 750, [255, 255, 255,
255]);

Screen('DrawText', window, 'When you are ready to start this procedure,
click the LEFT mouse button...', 50, 800, [255, 255, 255, 255]);

Screen(window, 'Flip');

buttons = 0; % When the user clicks the mouse, 'buttons' becomes
nonzero.
while buttons == 0
    [mX, mY, buttons] = GetMouse; % Check for a response...
    %T_Position = randperm(3,1);
```

```
end;


% Flip to the screen
Screen('Flip', window);


%% Initial blindspot localiser routine...
BS_XYs = [];

% Set Dot Translation Speed
Dot_Speed_per_sec = width/15;

% Loop the animation until a key is pressed
%while ~KbCheck
[mX,mY,buttons]=GetMouse;
while sum(buttons)~= 0
    [mX,mY,buttons]=GetMouse;
end

Preliminary_BS_Coords = zeros(1,4);
First_Guess_X = 0;
for BS_Est = 1 : 12

    BS_Est_Num = 1+mod(BS_Est-1,4);

    Repeat = 0;

    while Repeat == 0


        %% Sequence to initiate trial...
        SetMouse(width/2,height/2);
        [mX,mY,buttons]=GetMouse;
        while sum(buttons)~= 0
            [mX,mY,buttons]=GetMouse;
        end
        stop = 0;
        StartX=0;
        StartY=0;
        NowY=0;
        NowX=0;

        count=0;

        rand_pos1 = rand(1);
        if rand_pos1 < 0.5
            Y2 = Y1-Pixels_per_DVA;
        else
            Y2 = Y1+Pixels_per_DVA;
        end
        pause(0.1)

        while stop == 0
            [mX,mY,buttons]=GetMouse;

            if  Y1-(Y2-(StartY-NowY)) < -6 ||  Y1-(Y2-(StartY-NowY)) >
6
```

```
                Colour = [255, 0, 0, 0];
            else
                Colour = [0, 255, 0, 0];
                stop=1;
            end

            if sum(buttons) > 0 && count == 0
                StartX = mX;
                StartY = mY;
                NowX = mX;
                NowY=mY;
                count=count+1;
            elseif sum(buttons) > 0
                NowX=mX;
                NowY=mY;
            end
            Screen('DrawDots', window, [X1-(StartX-NowX) Y2-(StartY-
NowY)] , dot1SizePix, Colour, [], 2);
            Screen('FrameOval', window, Colour, [X1-Pixels_per_DVA/4
Y1-Pixels_per_DVA/4 X1+Pixels_per_DVA/4 Y1+Pixels_per_DVA/4], 3, 3);
            Screen('Flip', window);
        end
        Screen('FillOval', window, Colour, [100-Pixels_per_DVA/4
(height/3)-Pixels_per_DVA/4 100+Pixels_per_DVA/4
(height/3)+Pixels_per_DVA/4]);
        Screen('Flip', window);
        [mX,mY,buttons]=GetMouse;
        while sum(buttons)~= 0
            [mX,mY,buttons]=GetMouse;
        end


        %% Preliminary BS coordinate estimate procedure
        time = 0;
        tstrat = tic;
        while sum(buttons)== 0

            switch BS_Est_Num
                case 1
                    % Testing RIGHT eye
                    Xpos = X1 + (Dot_Speed_per_sec*time);
                    %Xpos = Xpos1;
                    Ypos = Y1+100;
                case 2
                    % Testing RIGHT eye
                    Xpos = scrnWidthPix-dot1SizePix -
(Dot_Speed_per_sec*time);
                    %Xpos = Xpos2;
                    Ypos = Y1+100;
                case 3
                    Xpos = First_Guess_X/2; %(Xpos1+Xpos2)/2;
                    Ypos = dot1SizePix + (Dot_Speed_per_sec*time);
                case 4
                    %Xpos = 0;
                    Xpos = First_Guess_X/2; %(Xpos1+Xpos2)/2;
                    Ypos = (height*0.75-dot1SizePix) -
(Dot_Speed_per_sec*time);
```

```
            end

            Screen('FillOval', window, Colour, [100-Pixels_per_DVA/4
(height/3)-Pixels_per_DVA/4 100+Pixels_per_DVA/4
(height/3)+Pixels_per_DVA/4]);
            Screen('FillOval', window, [0, 255, 0, 0], [Xpos-
dot1SizePix/2, Ypos-dot1SizePix/2, Xpos+dot1SizePix/2,
Ypos+dot1SizePix/2] );

            % Flip to the screen
            Screen(window, 'Flip');

            % Increment the time
            time = toc(tstrat);

            [mX,mY,buttons]=GetMouse;

        end

        while sum(buttons)>0
            [mX,mY,buttons]=GetMouse;
        end

        %% Check Response...
        Screen('TextFont',window, 'Courier New');
        Screen('TextStyle', window, 1+2);
        Screen('TextSize',window, 16);
        Screen('DrawText', window, 'Did you click as soon as the disc
dissapeared, but not before (Left Click)', 50, Y1-25, [255, 255, 255,
255]);
        Screen('DrawText', window, 'Or did you click early or late
(Right Click)', 50, Y1+25, [255, 255, 255, 255]);
        % Flip to the screen
        Screen('Flip', window);

        % buttons = [0 0 0]
        while sum(buttons)<1
            [mX,mY,buttons]=GetMouse;
        end
        if buttons(1) == 1
            Repeat = 1;
        end
        Screen('Flip', window);


    end

    if BS_Est <= 2
        First_Guess_X = First_Guess_X+Xpos;
    elseif BS_Est > 4
        if BS_Est_Num < 3
            Preliminary_BS_Coords(BS_Est_Num) =
Preliminary_BS_Coords(BS_Est_Num)+Xpos;
        else
            Preliminary_BS_Coords(BS_Est_Num) =
Preliminary_BS_Coords(BS_Est_Num)+Ypos;
        end
    end
```

```
end

% X and Y coordinates for Left'
BS_XYs(1,1) = Preliminary_BS_Coords(1)/2;

% X and Y coordinates for Right'
BS_XYs(2,1) = Preliminary_BS_Coords(2)/2;

% X and Y coordinates for Top'
BS_XYs(1,2) = Preliminary_BS_Coords(3)/2;

% X and Y coordinates for Bottom'
BS_XYs(2,2) = Preliminary_BS_Coords(4)/2;


BS_Centre = [BS_XYs(1,1)+((BS_XYs(2,1)-BS_XYs(1,1))/2)
BS_XYs(1,2)+((BS_XYs(2,2)-BS_XYs(1,2))/2)] ;
BS_Width = BS_XYs(2,1)-BS_XYs(1,1);
BS_Height = BS_XYs(2,2)-BS_XYs(1,2);


while sum(buttons)>0
    [mX,mY,buttons]=GetMouse;
end

%% Instructions...
Screen('TextFont',window, 'Courier New');
Screen('TextSize',window, 40);
Screen('TextStyle', window, 1+2);
Screen('DrawText', window, 'Instructions...', 0, 75, [255, 255, 255,
255]);

%Draw Text
Screen('TextSize',window, 16);
Screen('DrawText', window, 'In the next quick procedure you need to
report when moving discs APPEAR.', 50, 200, [255, 255, 255, 255]);
Screen('DrawText', window, 'Again, to begin each trial, you have to
slide a cursor into a small ring.', 50, 250, [255, 255, 255, 255]);

Screen('DrawText', window, 'Some time after the cursor turns green, a
moving disc will appear.', 50, 350, [255, 255, 255, 255]);
Screen('DrawText', window, 'Keep fixating the static green cursor, and
press the LEFT mouse button as soon as the moving disc', 50, 400, [255,
255, 255, 255]);
Screen('DrawText', window, 'appears. Reaction time is important, so
please press the mouse button as soon as the disc appears,', 50, 450,
[255, 255, 255, 255]);
Screen('DrawText', window, 'but NOT before!', 50, 500, [255, 255, 255,
255]);

Screen('DrawText', window, 'When you are ready to start this procedure,
click the LEFT mouse button...', 50, 660, [255, 255, 255, 255]);

Screen(window, 'Flip');

while sum(buttons)<1
    [mX,mY,buttons]=GetMouse;
```

```
end


% Flip to the screen
Screen('Flip', window);


%% Initial blindspot localiser routine %% inside out moving dot...
BS_XYs2 = [];


% Loop the animation until a key is pressed
%while ~KbCheck
[mX,mY,buttons]=GetMouse;
while sum(buttons)~= 0
    [mX,mY,buttons]=GetMouse;
end

Preliminary_BS_Coords=[];
for BS_Est = 1 : 8

    BS_Est_Num = 1+mod(BS_Est-1,4);

    Repeat = 0;
    while Repeat == 0


        if BS_Est_Num == 1
            Ypos = Y1+100; %yCenter; %yaha change kiya
        elseif BS_Est_Num == 2
            % Testing RIGHT eye
            tX = scrnWidthPix-dot1SizePix;
        elseif BS_Est_Num == 3
            'X and Y coordinates for Top'
            BS_XYs(1,2) = Ypos;
        else
            'X and Y coordinates for Bottom'
            BS_XYs(2,2) = Ypos;
        end


        %% Sequence to initiate trial...
        SetMouse(width/2,height/2);
        [mX,mY,buttons]=GetMouse;
        while sum(buttons)~= 0
            [mX,mY,buttons]=GetMouse;
        end
        stop = 0;
        StartX=0;
        StartY=0;
        NowY=0;
        NowX=0;

        count=0;

        rand_pos1 = rand(1);
        if rand_pos1 < 0.5
            Y2 = Y1-Pixels_per_DVA; %yaha change kiya hai
```

```
        else
            Y2 = Y1+Pixels_per_DVA; % yaha change kiya hai
        end
        pause(0.1)

        while stop == 0
            [mX,mY,buttons]=GetMouse;

            if  Y1-(Y2-(StartY-NowY)) < -6 ||  Y1-(Y2-(StartY-NowY)) >
6
                Colour = [255, 0, 0, 0];
            else
                Colour = [0, 255, 0, 0];
                stop=1;
            end

            if sum(buttons) > 0 && count == 0
                StartX = mX;
                StartY = mY;
                NowX = mX;
                NowY=mY;
                count=count+1;
            elseif sum(buttons) > 0
                NowX=mX;
                NowY=mY;
            end
            Screen('DrawDots', window, [X1-(StartX-NowX) Y2-(StartY-
NowY)] , dot1SizePix, Colour, [], 2);
            Screen('FrameOval', window, Colour, [X1-Pixels_per_DVA/4
Y1-Pixels_per_DVA/4 X1+Pixels_per_DVA/4 Y1+Pixels_per_DVA/4], 3, 3);
            Screen('Flip', window);
        end
        Screen('FillOval', window, Colour, [100-Pixels_per_DVA/4
(height/3)-Pixels_per_DVA/4 100+Pixels_per_DVA/4
(height/3)+Pixels_per_DVA/4]);
        Screen('Flip', window);
        [mX,mY,buttons]=GetMouse;
        while sum(buttons)~= 0
            [mX,mY,buttons]=GetMouse;
        end


        %% Preliminary BS coordinate estimate procedure
        X2 = BS_Centre(1);
        Y2 = BS_Centre(2);
        tX2 = X2-dot1SizePix;


        time = 0;
        tstrat = tic;
        while sum(buttons)== 0

            switch BS_Est_Num
                case 1
                    % Testing RIGHT eye
                    Xpos2 = X2 + (Dot_Speed_per_sec*time);
                    Ypos = BS_Centre(2);
```

```
                case 2
                    % Testing RIGHT eye
                    Xpos2 = tX2 - (Dot_Speed_per_sec*time);
                    Ypos = BS_Centre(2);

                case 3
                    Xpos2 = BS_Centre(1);
                    Ypos = (Y2-dot1SizePix) + (Dot_Speed_per_sec*time);

                case 4
                    Xpos2 = BS_Centre(1);
                    Ypos = (Y2-dot1SizePix) - (Dot_Speed_per_sec*time);

            end


            Screen('FillOval', window, Colour, [100-Pixels_per_DVA/4
(height/3)-Pixels_per_DVA/4 100+Pixels_per_DVA/4
(height/3)+Pixels_per_DVA/4]);
            Screen('DrawDots', window, [X2 Y2] , dot1SizePix, [255 255
255], [], 2);
            Screen('FillOval', window, [0, 255, 0, 0], [Xpos2-
dot1SizePix/2, Ypos-dot1SizePix/2, Xpos2+dot1SizePix/2,
Ypos+dot1SizePix/2] );

            % Flip to the screen
            Screen(window, 'Flip');

            % Increment the time
            time = toc(tstrat);

            [mX,mY,buttons]=GetMouse;

        end

        while sum(buttons)>0
            [mX,mY,buttons]=GetMouse;
        end

        %% Check Response...
        Screen('TextFont',window, 'Courier New');
        Screen('TextStyle', window, 1+2);
        Screen('TextSize',window, 16);
        Screen('DrawText', window, 'Did you click as soon as the disc
appeared, but not before (Left Click)', 50, Y1-25, [255, 255, 255,
255]);
        Screen('DrawText', window, 'Or did you click early or late
(Right Click)', 50, Y1+25, [255, 255, 255, 255]);
        % Flip to the screen
        Screen('Flip', window);

        % buttons = [0 0 0]
        while sum(buttons)<1
            [mX,mY,buttons]=GetMouse;
        end
        if buttons(1) == 1
            Repeat = 1;
        end
```

```
        Screen('Flip', window);

    end

    if BS_Est > 4
        if BS_Est_Num == 1
            'X and Y coordinates for Left'
            BS_XYs2(1,1) = Xpos2;
        elseif BS_Est_Num == 2
            'X and Y coordinates for Right'
            BS_XYs2(2,1) = Xpos2;
        elseif BS_Est_Num == 3
            'X and Y coordinates for Top'
            BS_XYs2(1,2) = Ypos;
        else
            'X and Y coordinates for Bottom'
            BS_XYs2(2,2) = Ypos;
        end
    end

end


BS_XYs_Avg(1,1)=(BS_XYs2(1,1)+BS_XYs(1,1))/2;  %Right
BS_XYs_Avg(2,1)=(BS_XYs2(2,1)+BS_XYs(2,1))/2;  %Left
BS_XYs_Avg(1,2)=(BS_XYs2(1,2)+BS_XYs(1,2))/2;  %Bottom
BS_XYs_Avg(2,2)=(BS_XYs2(2,2)+BS_XYs(2,2))/2;  %Top

BS_Centre2 = [BS_XYs_Avg(1,1)+((BS_XYs_Avg(2,1)-BS_XYs_Avg(1,1))/2)
BS_XYs_Avg(1,2)+((BS_XYs_Avg(2,2)-BS_XYs_Avg(1,2))/2)] ;
BS_Width2 = BS_XYs_Avg(1,1)-BS_XYs_Avg(2,1);
BS_Height2 = BS_XYs_Avg(1,2)-BS_XYs_Avg(2,2);


while sum(buttons)>0
    [mX,mY,buttons]=GetMouse;
end

%% Instructions for Fill-in...

Screen('TextFont',window, 'Courier New');
Screen('TextSize',window, 40);
Screen('TextStyle', window, 1+2);
Screen('DrawText', window, 'Instructions...', 0, 75, [255, 255, 255,
255]);

%Draw Text
Screen('TextSize',window, 16);
Screen('DrawText', window, 'This procedure will measure the position of
your physiological blindspot more precisely.', 50, 250, [255, 255, 255,
255]);
Screen('DrawText', window, 'To begin each trial, again you have to
slide a cursor into a small ring, and it will turn green.', 50, 350,
[255, 255, 255, 255]);

Screen('DrawText', window, 'Once the cursor turns green, keep looking
at it.', 50, 450, [255, 255, 255, 255]);
```

```
Screen('DrawText', window, 'Then there will be two green bars. At first
these will be clearly seperated. Your task is make them', 50, 500,
[255, 255, 255, 255]);
Screen('DrawText', window, 'stretch (by pressing the LEFT mouse button)
until they seem to touch. As soon as they TOUCH, but', 50, 550, [255,
255, 255, 255]);
Screen('DrawText', window, 'NOT BEFORE, release the LEFT mouse button
(fast response times are important). Then you should press', 50, 600,
[255, 255, 255, 255]);
Screen('DrawText', window, 'the RIGHT mouse button, and the next trial
will begin...', 50, 650, [255, 255, 255, 255]);


Screen('DrawText', window, 'When you are ready to start this procedure,
click the LEFT mouse button...', 50, 750, [255, 255, 255, 255]);

Screen(window, 'Flip');

while sum(buttons)<1
    [mX,mY,buttons]=GetMouse;
end


% Flip to the screen
Screen('Flip', window);

%% Initial fill-in ...


Element_Wd = 2;
Element_Hgt = 0.5;

Animation_Duration = 1;
Start_Distance = 3;
Travel_Distance = 5;

Num_Trials = 10;


%%
BS_Coords = BS_Centre2;
BS_Coords(3) = BS_Width2;
BS_Coords(4) = BS_Height2;



for count = 1:Num_Trials


    %% Sequence to initiate trial...
    SetMouse(width/2,height/2);
    [mX,mY,buttons]=GetMouse;
    while sum(buttons)~= 0
        [mX,mY,buttons]=GetMouse;
    end
    stop = 0;
    StartX=0;
    StartY=0;
```

```
    NowY=0;
    NowX=0;

    init_count=0;

    rand_pos1 = rand(1);
    if rand_pos1 < 0.5
        Y2 = Y1-Pixels_per_DVA;
    else
        Y2 = Y1+Pixels_per_DVA;
    end
    pause(0.1)

    %[Start_MouseX,Start_MouseY,buttons]=GetMouse;
    while stop == 0
        [mX,mY,buttons]=GetMouse;

        if  Y1-(Y2-(StartY-NowY)) < -6 ||  Y1-(Y2-(StartY-NowY)) > 6
            Colour = [255, 0, 0, 0];
        else
            Colour = [0, 255, 0, 0];
            stop=1;
        end

        if sum(buttons) > 0 && init_count == 0
            StartX = mX;
            StartY = mY;
            NowX = mX;
            NowY=mY;
            init_count=init_count+1;
        elseif sum(buttons) > 0
            NowX=mX;
            NowY=mY;
        end
        %NowX = Start_MouseX-mX;
        Screen('DrawDots', window, [X1-(StartX-NowX) Y2-(StartY-NowY)]
, dot1SizePix, Colour, [], 2);
        %Screen('DrawDots', windowPtr, xy [,size] [,color] [,center]
[,dot_type][,lenient]);
        Screen('FrameOval', window, Colour, [X1-Pixels_per_DVA/4 Y1-
Pixels_per_DVA/4 X1+Pixels_per_DVA/4 Y1+Pixels_per_DVA/4], 3, 3);
        Screen('Flip', window);
    end
        Screen('FillOval', window, Colour, [100-Pixels_per_DVA/4
(height/3)-Pixels_per_DVA/4 100+Pixels_per_DVA/4
(height/3)+Pixels_per_DVA/4]);
        Screen('Flip', window);


 %% Test protocol...
    BS_Colour = [255, 0, 0, 0];
    Screen('FillOval', window, Colour, [100-Pixels_per_DVA/4
(height/3)-Pixels_per_DVA/4 100+Pixels_per_DVA/4
(height/3)+Pixels_per_DVA/4]);

    SetMouse(width/2,height/2);
    [mX,mY,buttons]=GetMouse;
    while sum(buttons)~= 0
```

```
        [mX,mY,buttons]=GetMouse;
    end

    stop = 0;



    BS_Colour = [255, 0, 0, 0];
    Stop = 0;
    Wd = 11.75;
    while Stop == 0
        [mX,mY,buttons]=GetMouse;
        Screen('FillOval', window, Colour, [100-Pixels_per_DVA/4
(height/3)-Pixels_per_DVA/4 100+Pixels_per_DVA/4
(height/3)+Pixels_per_DVA/4]);
        Screen('FillRect', window, [0 255 0], [BS_Coords(1)-
(6*Pixels_per_DVA) BS_Coords(2)-(Element_Hgt/2)*Pixels_per_DVA
BS_Coords(1)+(6*Pixels_per_DVA)
BS_Coords(2)+(Element_Hgt/2)*Pixels_per_DVA]);
        Screen('FillRect', window, [0 0 0], [BS_Coords(1)-
(Wd/2*Pixels_per_DVA) BS_Coords(2)-(Element_Hgt/2)*Pixels_per_DVA
BS_Coords(1)+(Wd/2*Pixels_per_DVA)
BS_Coords(2)+(Element_Hgt/2)*Pixels_per_DVA]);

        if buttons(3) == 1
            Stop = 1;
        elseif buttons(1) == 1
            Wd = Wd-0.05;
        elseif buttons(2) == 1
            Wd = Wd+0.05;
        end

        if Wd > 11.75
            Wd = 11.75;
        elseif Wd < 1
            Wd = 1;
        end
        Screen('Flip', window);
    end


    [mX,mY,buttons]=GetMouse;
    while sum(buttons)~= 0
        [mX,mY,buttons]=GetMouse;
    end

    Results_Array(count) = Wd;

    fprintf(fid, '\nWidth Setting %5.2f', Wd);

end

Mean_Wd = mean(Results_Array);


%% Instructions fLICKER ...

Screen('TextFont',window, 'Courier New');
```

```
Screen('TextSize',window, 40);
Screen('TextStyle', window, 1+2);
Screen('DrawText', window, 'Instructions...', 0, 75, [255, 255, 255,
255]);

%Draw Text
Screen('TextSize',window, 16);
Screen('DrawText', window, 'Now we are going to take a TIMING
measure.', 50, 150, [255, 255, 255, 255]);

Screen('DrawText', window, 'Again, to begin each trial you have to
slide the cursor into the ring.', 50, 200, [255, 255, 255, 255]);

Screen('DrawText', window, 'When the cursor turns green, two flickering
squares will appear', 50, 250, [255, 255, 255, 255]);
Screen('DrawText', window, 'DONT LOOK AT THEM! Keep fixating the static
green cursor.', 50, 300, [255, 255, 255, 255]);

Screen('DrawText', window, 'Your task is to say if the squares were
flickering on and off together', 50, 350, [255, 255, 255, 255]);
Screen('DrawText', window, 'or they were flickering at different times
- creating a sense of motion.', 50, 400, [255, 255, 255, 255]);

Screen('DrawText', window, 'Before you begin people ask NEHA to start
EEG recording.', 50, 500, [255, 255, 255, 255]);

Screen('DrawText', window, 'When you are ready to begin, click the LEFT
mouse button...', 50, 550, [255, 255, 255, 255]);

Screen(window, 'Flip');

buttons = 0; % When the user clicks the mouse, 'buttons' becomes
nonzero.
while buttons == 0
    [mX, mY, buttons] = GetMouse; % Check for a response...
    %T_Position = randperm(3,1);
end

% Flip to the screen
Screen('Flip', window);


Adjacent = BS_Coords(1)-100;
Opposite = BS_Coords(2)-(height/3);
Hypotenuse = sqrt((Adjacent*Adjacent)+(Opposite*Opposite));
Angle = atand(Opposite/Adjacent);

Control_Angle = Angle-60;

Control_X = cosd(Control_Angle)*Hypotenuse;
Control_Y = -(sind(Control_Angle)*Hypotenuse);

Check_Angle = atand(Control_X/Control_Y)-Control_Angle
Check_Hypotenuse = sqrt((Control_Y*Control_Y)+(Control_X*Control_X))-
Hypotenuse

%% Initial Temporal filling-in ...
```

```
SD_Results = zeros(3,4);

N_Repeats = 3;

Num_Trials = N_Repeats*7;
Trial_Order = randperm(Num_Trials);


% Get framerate
Hz = FrameRate(window);
BS_Rt_Side = BS_Coords(1)+(Mean_Wd/2*Pixels_per_DVA);
BS_Lt_Side = BS_Coords(1)-(Mean_Wd/2*Pixels_per_DVA);

Time = datestr(now);
fprintf(fid, '\n\nStarted Main Experiment: %s \n', Time);


Animation_Duration = 10;
for count = 1:Num_Trials
    count

    if Trial_Order(count)<= Num_Trials/6
        % Testing at blindspot - In-phase flicker
        Condition = 1;
        Results_Row = 1;
        Test_Y1 = BS_Coords(2);
        Test_Y2 = Test_Y1;
        Test_Lt_Side = BS_Coords(1)-((Mean_Wd/2)*Pixels_per_DVA);
        Test_Rt_Side = BS_Coords(1)+((Mean_Wd/2)*Pixels_per_DVA);
        if EEG_Session == 1
            invoke(RP,'SetTagVal','trig_val',Trigger_Vals(1)); % to
signal start of Blindspot Flicker Trial
        end
    elseif Trial_Order(count) > Num_Trials/6 & Trial_Order(count)<=
(Num_Trials/6)*2
        % Testing at blindspot - Apparent Motion
        Condition = 2;
        Results_Row = 1;
        Test_Y1 = BS_Coords(2);
        Test_Y2 = Test_Y1;
        Test_Lt_Side = BS_Coords(1)-(Mean_Wd/2*Pixels_per_DVA);
        Test_Rt_Side = BS_Coords(1)+(Mean_Wd/2*Pixels_per_DVA);
        if EEG_Session == 1
            invoke(RP,'SetTagVal','trig_val',Trigger_Vals(2)); % to
signal start of Blindspot Apparent Motion Trial
        end
    elseif Trial_Order(count) > (Num_Trials/6)*2 & Trial_Order(count)<=
(Num_Trials/6)*3
        % Testing at control position - In-phase flicker
        Condition = 3;
        Results_Row = 2;
        Test_Y1 = (height/3)+Control_Y;
        Test_Y2 = Test_Y1;
        Test_Lt_Side = Control_X+100-(Mean_Wd/2*Pixels_per_DVA);
        Test_Rt_Side = Control_X+100+(Mean_Wd/2*Pixels_per_DVA);
        if EEG_Session == 1
            invoke(RP,'SetTagVal','trig_val',Trigger_Vals(3)); % to
signal start of Control Flicker Trial
```

```
        end
    elseif Trial_Order(count) > (Num_Trials/6)*3 & Trial_Order(count)<=
(Num_Trials/6)*4
        % Testing at control position - Apparent Motion
        Condition = 4;
        Results_Row = 2;
        Test_Y1 = (height/3)+Control_Y;
        Test_Y2 = Test_Y1;
        Test_Lt_Side = Control_X+100-(Mean_Wd/2*Pixels_per_DVA);
        Test_Rt_Side = Control_X+100+(Mean_Wd/2*Pixels_per_DVA);
        if EEG_Session == 1
            invoke(RP,'SetTagVal','trig_val',Trigger_Vals(4)); % to
signal start of Control Apparent Motion Trial
        end
    elseif Trial_Order(count) > (Num_Trials/6)*4 & Trial_Order(count)<=
(Num_Trials/6)*5
        'Testing control to blindspot - In-phase flicker'
        Condition = 5;
        Results_Row = 3;
        Test_Y1 = BS_Coords(2);
        Test_Y2 = (height/3)+Control_Y;
        Position = rand(1);
        if Position < 0.5
            Test_Lt_Side = BS_Coords(1)-(Mean_Wd/2*Pixels_per_DVA);
            Test_Rt_Side = Control_X+100+(Mean_Wd/2*Pixels_per_DVA);
        else
            Test_Lt_Side = BS_Coords(1)+(Mean_Wd/2*Pixels_per_DVA);
            Test_Rt_Side = Control_X+100-(Mean_Wd/2*Pixels_per_DVA);
        end
        if EEG_Session == 1
            invoke(RP,'SetTagVal','trig_val',Trigger_Vals(1)); % to
signal start of Blindspot to Control Flicker Trial
        end
    else
        'Testing control to blindspot - Apparent Motion'
        Condition = 6;
        Results_Row = 3;
        Test_Y1 = BS_Coords(2);
        Test_Y2 = (height/3)+Control_Y;
        Position = rand(1);
        if Position < 0.5
            Test_Lt_Side = BS_Coords(1)-(Mean_Wd/2*Pixels_per_DVA);
            Test_Rt_Side = Control_X+100+(Mean_Wd/2*Pixels_per_DVA);
        else
            Test_Lt_Side = BS_Coords(1)+(Mean_Wd/2*Pixels_per_DVA);
            Test_Rt_Side = Control_X+100-(Mean_Wd/2*Pixels_per_DVA);
        end
        if EEG_Session == 1
            invoke(RP,'SetTagVal','trig_val',Trigger_Vals(2)); % to
signal start of Blindspot to Control Flicker Trial
        end
    end

    %% Sequence to initiate trial for Temporal filling In...

    SetMouse(width/2,height/2);
    [mX,mY,buttons]=GetMouse;
    while sum(buttons)~= 0
```

```
        [mX,mY,buttons]=GetMouse;
    end
    stop = 0;
    StartX=0;
    StartY=0;
    NowY=0;
    NowX=0;

    init_count=0;

    rand_pos1 = rand(1);
    if rand_pos1 < 0.5
        Y2 = (height/3)-Pixels_per_DVA;
    else
        Y2 = (height/3)+Pixels_per_DVA;
    end
    pause(0.1)

    while stop == 0
        [mX,mY,buttons]=GetMouse;

        if  (height/3)-(Y2-(StartY-NowY)) < -6 ||  (height/3)-(Y2-
(StartY-NowY)) > 6
            Colour = [255, 0, 0, 0];
        else
            Colour = [0, 255, 0, 0];
            stop=1;
        end

        if sum(buttons) > 0 && init_count == 0
            StartX = mX;
            StartY = mY;
            NowX = mX;
            NowY=mY;
            init_count=init_count+1;
        elseif sum(buttons) > 0
            NowX=mX;
            NowY=mY;
        end
        Screen('DrawDots', window, [100-(StartX-NowX) Y2-(StartY-NowY)]
, dot1SizePix, Colour, [], 2);
        Screen('FrameOval', window, Colour, [100-Pixels_per_DVA/4
(height/3)-Pixels_per_DVA/4 100+Pixels_per_DVA/4
(height/3)+Pixels_per_DVA/4], 3, 3);
        Screen('Flip', window);
    end
    Screen('FillOval', window, Colour, [100-Pixels_per_DVA/4
(height/3)-Pixels_per_DVA/4 100+Pixels_per_DVA/4
(height/3)+Pixels_per_DVA/4]);
    Screen('Flip', window);


    %% Test protocol...

    Element_Wd = 1.5;
    Element_Hgt = 0.5;

    Col1 = [255 255 255];
```

```
    Col2 = [0 0 0];
    if mod(Condition,2) == 1
        Col1b = [255 255 255];
        Col2b = [0 0 0];
    else
        Col1b = [0 0 0];
        Col2b = [255 255 255];
    end


    BS_Colour = [255, 0, 0, 0];
    Screen('DrawDots', window, [100-(StartX-NowX) Y2-(StartY-NowY)] ,
dot1SizePix, Colour, [], 2);

    if EEG_Session == 1 && Condition > 4
        vsg(vsgSetDisplayPage,1+ vsgTRIGGERPAGE); %Send Trigger
        pause(0.25);
        vsg(vsgSetDisplayPage,1+ vsgTRIGGERPAGE); %Send Trigger
    elseif EEG_Session == 1 && Condition <= 4
        vsg(vsgSetDisplayPage,1+ vsgTRIGGERPAGE); %Send Trigger
    end
    time = 0;
    tstrat = tic;
    while time < Animation_Duration

        time = toc(tstrat);
        if mod(time,1/4) <= 1/8
            ColA = Col1;
            ColB = Col1b;
        else
            ColA = Col2;
            ColB = Col2b;
        end

        Screen('DrawDots', window, [100-(StartX-NowX) Y2-(StartY-NowY)]
, dot1SizePix, Colour, [], 2);

%         Screen('FillRect', window, ColA, [BS_Rt_Side Test_Y1-
(2*Element_Hgt)*Pixels_per_DVA BS_Rt_Side+(Element_Wd*Pixels_per_DVA)
Test_Y1+(2*Element_Hgt)*Pixels_per_DVA]);
%         Screen('FillRect', window, ColB, [BS_Lt_Side-
(Element_Wd*Pixels_per_DVA) Test_Y2-(2*Element_Hgt)*Pixels_per_DVA
BS_Lt_Side Test_Y2+(2*Element_Hgt)*Pixels_per_DVA]);

        Screen('FillRect', window, ColB, [BS_Lt_Side-
(Element_Wd*Pixels_per_DVA) Test_Y2-(2*Element_Hgt/2)*Pixels_per_DVA
BS_Lt_Side Test_Y2+(2*Element_Hgt/2)*Pixels_per_DVA]);
        Screen('FillRect', window, ColA, [BS_Rt_Side Test_Y1-
(2*Element_Hgt/2)*Pixels_per_DVA BS_Rt_Side+(Element_Wd*Pixels_per_DVA)
Test_Y1+(2*Element_Hgt/2)*Pixels_per_DVA]);

        Screen('Flip', window);

    end
    Screen('Flip', window);


    %% Take Response...
```

```
    Screen('TextFont',window, 'Courier New');
    Screen('TextStyle', window, 1+2);
    Screen('TextSize',window, 16);
    Screen('DrawText', window, 'Were the squares flickering on and off
together (Left Click)', 50, Y1-25, [255, 255, 255, 255]);
    Screen('DrawText', window, 'Or at different times - creating a
sense of motion (Right Click)', 50, Y1+25, [255, 255, 255, 255]);
    % Flip to the screen

    SetMouse(width/2,height/2);
    [mX,mY,buttons]=GetMouse;
    while sum(buttons)~= 0
        [mX,mY,buttons]=GetMouse;
    end

    Screen('Flip', window);
    while sum(buttons)== 0
        [mX,mY,buttons]=GetMouse;
    end

    if buttons(1) == 1
        % Flicker Reported
        if mod(Condition,2) == 1
            % Flicker Presented - Correct Rejection
            SD_Results(Results_Row,4)=SD_Results(Results_Row,4)+1;
            Correct = 1;
            Option = 1;
        else
            % AM Presented - Miss
            SD_Results(Results_Row,2)=SD_Results(Results_Row,2)+1;
            Correct = 0;
            Option = 2;
        end
    elseif buttons(3) == 1
        % Apparent Motion Reported
        if mod(Condition,2) == 1
            % Flicker Presented - False Alarm
            SD_Results(Results_Row,3)=SD_Results(Results_Row,3)+1;
            Correct = 0;
            Option = 3;
        else
            % AM Presented - Hit
            SD_Results(Results_Row,1)=SD_Results(Results_Row,1)+1;
            Correct = 1;
            Option = 4;
        end
    end

    Screen('TextSize',window, 22);
    if Correct == 1
        Screen('DrawText', window, 'CORRECT :)', 200, Y2, [0, 255, 0]);
    else
        Screen('DrawText', window, 'INCORRECT :(', 200, Y2, [255, 0,
0]);
    end
    Screen('Flip', window);
```

```
    fprintf(fid, '\nTrialCondition %5.0f \tSelection %5.0f' ,
Condition, Option);


    pause(0.4)


end

mu = 0;
sigma = 1;


fprintf(fid, '\n\n\n\nRESULT SUMMARIES');

fprintf(fid, '\n\nBlindspot distance from fixation: %5.1f
(dva)',(Hypotenuse / Pixels_per_DVA) );
fprintf(fid, '\n\nBlindspot X-distance from fixation: %5.1f
(dva)',(BS_Coords(1)-X1) / Pixels_per_DVA);
fprintf(fid, '\nBlindspot Y-distance from fixation: %5.1f
(dva)',(BS_Coords(2)-Y1) / Pixels_per_DVA_height);
fprintf(fid, '\nBlindspot Width: %5.1f (dva)',BS_Coords(3) /
Pixels_per_DVA);
fprintf(fid, '\nBlindspot Height: %5.1f (dva)',BS_Coords(4) /
Pixels_per_DVA_height);


fprintf(fid, '\n\n\nBlindspot Trials:');
fprintf(fid, '\nHits \tMisses \tFAs \tCRs');
fprintf(fid, '\n%4.0f \t\t\t%4.0f \t\t%4.4f \t\t%4.0f',
SD_Results(1,1),SD_Results(1,2),SD_Results(1,3),SD_Results(1,4));
HR = SD_Results(1,1) / (SD_Results(1,1)+SD_Results(1,2));
FAR = SD_Results(1,3)/ (SD_Results(1,3)+SD_Results(1,4));
InvHR = norminv(HR,mu,sigma);
InvFAR = norminv(FAR,mu,sigma);
dprime = InvHR - InvFAR;
c = -(InvHR + InvFAR) / 2;
fprintf(fid, '\n\nHit Rate: %4.3f \nFAR: %4.3f \ndPrime: %4.3f \nCrit:
%4.3f',HR,FAR,dprime,FAR);

fprintf(fid, '\n\n\nControl Position Trials:');
fprintf(fid, '\nHits \tMisses \tFAs \tCRs');
fprintf(fid, '\n%4.0f \t\t\t%4.0f \t\t%4.4f \t\t%4.0f',
SD_Results(2,1),SD_Results(2,2),SD_Results(2,3),SD_Results(2,4));
HR = SD_Results(2,1) / (SD_Results(2,1)+SD_Results(2,2));
FAR = SD_Results(2,3)/ (SD_Results(2,3)+SD_Results(2,4));
InvHR = norminv(HR,mu,sigma);
InvFAR = norminv(FAR,mu,sigma);
dprime = InvHR - InvFAR;
c = -(InvHR + InvFAR) / 2;
fprintf(fid, '\n\nHit Rate: %4.3f \nFAR: %4.3f \ndPrime: %4.3f \nCrit:
%4.3f',HR,FAR,dprime,FAR);
fprintf(fid, '\n\n');


fprintf(fid, '\n\n\nControl to Blindspot Position Trials:');
fprintf(fid, '\nHits \tMisses \tFAs \tCRs');
```

```
fprintf(fid, '\n%4.0f \t\t\t%4.0f \t\t%4.4f \t\t%4.0f',
SD_Results(3,1),SD_Results(3,2),SD_Results(3,3),SD_Results(3,4));
HR = SD_Results(2,1) / (SD_Results(2,1)+SD_Results(2,2));
FAR = SD_Results(2,3)/ (SD_Results(2,3)+SD_Results(2,4));
InvHR = norminv(HR,mu,sigma);
InvFAR = norminv(FAR,mu,sigma);
dprime = InvHR - InvFAR;
c = -(InvHR + InvFAR) / 2;
fprintf(fid, '\n\nHit Rate: %4.3f \nFAR: %4.3f \ndPrime: %4.3f \nCrit:
%4.3f',HR,FAR,dprime,FAR);
fprintf(fid, '\n\n');

Time = datestr(now);
fprintf(fid, '\n\nFinished: %s \n', Time);

status = fclose('all');
ShowCursor
sca
```
